# Benchmarking Computational Methods for Estimating the Pathogenicity of Wilson’s Disease Mutations

**DOI:** 10.1101/780924

**Authors:** Ning Tang, Thomas Sandahl, Peter Ott, Kasper P. Kepp

## Abstract

Genetic variations in the gene encoding the copper-transport protein ATP7B are the primary cause of Wilson’s disease. Controversially, clinical prevalence seems much smaller than prevalence estimated by genetic screening tools, causing fear that many people are undiagnosed although early diagnosis and treatment is essential. To address this issue, we benchmarked 16 state-of-the-art computational disease-prediction methods against established data of missense *ATP7B* mutations. Our results show that the quality of the methods vary widely. We show the importance of optimizing the threshold of the methods used to distinguish pathogenic from non-pathogenic mutations against data of clinically confirmed pathogenic and non-pathogenic mutations. We find that most methods use thresholds that predict too many ATP7B mutations to be pathogenic. Thus, our findings explain the current controversy on Wilson’s disease prevalence, because meta analysis and text search methods include many computational estimates that lead to higher disease prevalence than clinically observed. Since proteins differ widely, a one-size-fits-all threshold for all proteins cannot distinguish efficiently pathogenic and non-pathogenic mutations, as shown here. We also show that amino acid changes with small evolutionary substitution probability, mainly due to amino acid volume, are more associated with disease, implying a pathological effect on the conformational state of the protein, which could affect copper transport or ATP recognition and hydrolysis. These findings may be a first step towards a more quantitative genotype-phenotype relationship of Wilson’s disease.

## Introduction

Wilson’s disease (WD) is a rare autosomal recessive disorder of copper metabolism caused by pathogenic variants of the human *ATP7B* gene encoding the ATP7B protein, which is a copper-transporting P-type ATPase.^1, 2^ The approximately 160-kDa membrane protein contains a large N-terminal domain consisting of six metal-binding domains, eight transmembrane segments (TMs), an adenosine triphosphate (ATP) binding domain, and a soluble C-terminal tail.^3–8^ In the hepatocyte, ATP7B transports copper from the cytosol into the Golgi apparatus and mediates either the incorporation of copper into ceruloplasmin or the excretion of excess copper into the bile.^9, 10^ To accomplish its copper transport function, the protein depends critically on its ability to use the energy gained by ATP hydrolysis.^3^

Pathogenic *ATP7B* mutations cause loss of copper transporting function resulting in accumulation of copper in multiple organs, most notably brain, liver, and kidney^11–13^ As a result, WD patients present with either hepatic, neurologic or psychiatric symptoms, often in combination.^14, 15^ If left untreated, WD with chronic presentation is fatal within a 5−10 year period from first symptom onset. However, apart from the rare acute fulminant hepatic presentation that requires liver transplantation, proper medical treatment can typically ensure a near-normal life expectancy, but this depends critically on early and accurate diagnosis.^18–21^

The clinical handling of patients with WD faces two major challenges; the uncertainty of the prevalence of the disease (the number of affected people within a given population) and the lack of a clear genotype-phenotype relation that enables the estimate of disease severity and manifestation. The worldwide prevalence was estimated in 1984 to be around 1 in 30,000^22^, corresponding to a carrier frequency of approximately 1 in 90, considered a reasonable estimate.^18, 23, 24^ Due to the heterogeneity in the clinical presentation and the age of presentation, a substantial number of patients are undiagnosed.^25^ Prevalence approaches 1:30,000 in some countries where diagnostic awareness of WD is high, such as Austria^26^ and France^27^. However, recent population-genetic studies based on the computer analysis of observed variants have led to estimates of WD prevalence of 1:1,400^28^, 1:7,100^29^ or 1:6,500^30^. If true, these studies suggest that at least 75% of people affected by the disease are undiagnosed with potentially fatal consequences.

However, the difference between clinical observations and genetic predictions may also raise questions regarding the validity of current computational analysis of observed genetic variants, which are widely used to estimate pathogenicity of genetic variants. Computational analysis of pathogenicity of mutations is increasingly used to examine disease mechanisms and to categorize pathogenic and non-pathogenic variants.^31^ Some computational methods have shown ability to identify pathogenic mutations.^31–33^ These predictive methods use the evolutionary “unlikeliness” of the amino acid substitution, the involved change in biochemical properties, and 3-dimensional protein structure to classify mutations. Loss of protein stability potentially leading to partial loss of function is a common feature of many inherited diseases. In such cases, structure-based computational methods can identify pathogenic mutations.^34, 35^ Alternatively, evolutionary conservation information may capture very disruptive amino acid changes that are unlikely to occur during natural evolution typically because they impair protein conformational integrity or function.^36^

We note that from a molecular evolution perspective, it is expected that a majority of naturally occurring protein variants are nearly neutral in their functional effect, which is the basis for the so-called neutral theory of evolution and the widely supported use of molecular clocks in phylogeny^37^. In the clinical terminology^38^, these probably tend to be the benign variants. This insight is further complicated by penetrance being modulated by non-genetic and genetic confounding factors. We hypothesized that, since proteins differ widely in size, shape, stability, location, and natural function, the impact of a typical human mutation may be very protein-dependent. For example, abundant proteins are known to evolve much more slowly than less abundant proteins due to selection pressures, and evolution rate is also very dependent on specific selection pressures of the protein^39–42^. Yet most methods suggest a default threshold to distinguish pathogenic from non-pathogenic mutations. From a clinical strategic perspective there is an urgent need to solve this issue and identify methods that correctly distinguish truly disease-causing mutations from benign (neutral) variants^38^, and possibly also the severity and penetrance of the pathogenic mutations.

As ATP7B is the only identified gene known to cause WD, genetic screening for known pathogenic variants is a sensitive approach to diagnose WD. However, in cases where the functional impact of a variant is unclear, genetic testing only provides circumstantial evidence, as variants display diverse functional effects. Further complications such as life-style and environmental risk modifiers and the low frequency and unknown penetrance of the mutations complicate diagnosis even further.^43^ Direct functional testing of disease-causing ATP7B mutations would ideally be the most sensitive method to diagnose WD, but such functional tests of each new variant is time-consuming. Clarification of these issues may affect the use of genetic testing for diagnosing patients suspected of having WD^18^ and may also have implications for conclusions based on genetic population screenings.

Genotype-phenotype relations would aid our understanding of the pathophysiology and the development of new diagnostic and therapeutic strategies. Such relations have so far met with little success.^19^ More than 700 ATP7B natural variants have been identified, including mostly missense mutations, insertions/deletions, and some rare splice-site mutations as summarized in the WD database (http://www.wilsondisease.med.ualberta.ca).^44^ Whereas truncating mutations tend to severely impair copper metabolism and cause early age of disease onset, some mutations do not relate to hepatic or neurologic presentations, supporting the concept of a pool of disease-wise benign natural ATP7B variants.^45, 46^ For missense mutations the genotype-phenotype relationship is even weaker as they cause a wide variety of symptoms in WD patients implying that they may affect ATP7B function in different ways.^47, 48^ These missense mutations are distributed across the ATP7B gene, but tend to cluster in the ATP binding domain indicating its importance for the ATP-dependent copper transport function.^49^ There is considerable phenotypic variation between individuals with the same mutation, even within the same families and in monozygotic twins^50, 51^ showing clearly the need for understanding the risk modulation effects of specific mutations on ATP7B functionality and clinical presentation.

In the present study we performed a detailed computational study of ATP7B protein variants using 16 widely used structure- and sequence-based methods with the specific aims i) to test the application of state of the art computational screening methods to the problem of WD where diagnosis is challenged; ii) to identify amino acid properties that correlate with disease presentation. We show that several sequence-based methods can accurately classify pathogenic variants if the threshold is optimized before quantitative diagnosis of WD. However, outcomes are extremely dependent on the thresholds used and default thresholds overestimate disease prevalence because of the presence of nearly neutral (benign) variants. This finding can largely explain the discrepancy between genetic-screening based and clinically observed WD prevalence. We also identify several important chemical features that determine whether a variant is disease-causing or not, which may aid the so far unsuccessful understanding of WD genotype-phenotype relations. In particular, the molecular volume of the amino acids is the only simple amino acid property that predicts disease tendency among 48 studied properties, pointing towards an effect on membrane conformational integrity that may aid an understanding of the molecular mechanism of WD pathology.

## Computational Methods

### Data for ATP7B genetic variants

We studied several data sets of ATB7B variants related to WD, but ultimately settled on using the mutations from the WD database^44^ (http://www.wilsondisease.med.ualberta.ca) for reasons described below. As most of the studied methods can only deal with missense mutations, only these were selected for investigation, which substantially reduces the number of relevant data points and affects the data set choice. There are two main concerns: 1) the completeness of the dataset and 2) the confidence in the assignment of the clinical impact of each variant.

When comparing to the most recent 2019 database by Gao et al.^29^ we found that the WD database includes almost all variants with high confidence of pathogenicity according to the more recent criteria by Richards et al.^38^ We tested the sensitivity of our conclusions by including the most recent variants from 2019.^29^ As shown below, this did not affect our conclusions, mainly because the confidently assigned variants have changed little compared to the major increase in the total number of inferred variants from genome screening, since functional and clinical testing has not experienced the same growth in capacity as sequencing and computational screening tools.

Many of the most confidently assigned loss-of-function variants are not missense mutations (e.g. deletions) and not studied by the applied methods; our analysis deals mainly with the difficult grey zone of missense mutations that are commonly nearly neutral (benign) and are the cause of the current controversy on disease prevalence. Gao et al.^29^ use broad screening approaches (including text search and meta analysis) to maximize completeness at the expense of confidence in the assignment of pathogenicity. The new data thus contain many computational estimates of pathogenicity; these estimates emerged mainly during the last decade and in the case of WD, after the WD database was complete. We find that the WD database, by minimizing recent computational and low-confidence screening results, is optimal for the analysis that we conduct here, where we specifically want to avoid pollution by computational screening estimates in the benchmark data. The insensitivity of our findings to reasonable variations in data set are discussed later in this paper.

The WD dataset includes 722 and 172 entries for pathogenic and non-pathogenic variants, respectively; of these, 291 variants are unique missense mutations studied in the present work. Clinical effects from loss of function studies (class 2) provide the best evidence for these mutations^44^. Since WD is a loss-of-function disease, functional studies provide a strong support for pathogenicity in particular in combination with control data for normal people (class 4). We estimate that, compared to the classification by Richards et al.^38^, which was not available and thus not used in the WD database, the loss-of-function feature makes the confidence of pathogenicity approximately strong (PS), which is the best possible situation as statistical critical mass of data points is still required. Among the 291 missense mutations studied, there are 267 pathogenic mutations and 24 non-pathogenic mutations. In addition, 15 phenotypic mutations were reported as both pathogenic and non-pathogenic; these mutations were not included in our dataset as their pathogenicity is variable or debated. Details of the used mutation data set are shown in **Table S1**.

### Studying mutations by computational mutagenesis using structure-based methods

Since the full ATP7B protein structure is not available, homology models will be unreliable. Instead, the available NMR structures of several domains were used to perform structure-based mutation analysis where possible. As shown previously^32, 52–54^ the ΔΔG (the change in stability caused by point mutation) is very structure-dependent for some stability-prediction methods but less so for others, and this means that more methods and structures should be compared whenever possible. Five structures were used with the Protein Data Bank (PDB) IDs 2N7Y, 2LQB, 2ROP, 2EW9 and 2ARF corresponding to metal-binding domains 1 (2N7Y), 2 (2LQB), 3 and 4 (2ROP), 5 and 6 (2EW9), and the ATP binding domain (2ARF), respectively.^55–59^

For FoldX (version 5),^60^ the structures were first repaired using the RepairPDB function, and then the BuildModel function was applied to the repaired structures to obtain ΔΔG values from five independent runs. The final reported ΔΔG values were the averages of these five independent runs for each mutation. For Rosetta (2019.07.60616 weekly release version), the structures were relaxed to produce 20 optimized structures. The structure with the lowest score was then used for ΔΔG calculation using the Cartesian version of Rosetta DDG protocol with three iterations.^61, 62^ The final ΔΔG values were calculated based on the difference in total scores averaged over three rounds for the mutant and wild type structures. For I-mutant (version 2.0),^63^ the secondary structure of the different domains was calculated using the DSSP algorithm.^64^ The obtained results and the corresponding structures were then submitted to I-mutant for calculating the protein stability changes upon single site mutation. For mCSM,^65^ SDM,^66^ and DUET,^67^ the web server versions of the programs were used with default settings for calculating the ΔΔG values using the original NMR ensemble as input. Four mutations (S291A, S291I, S291Q and M573H) could not be computed by SDM and DUET and were thus not included in the final SDM and DUET results. For POPMUSIC,^68^ HOTMUSIC,^69^ and SNPMUSIC,^70^ the calculations were performed using the DEZYME (http://www.dezyme.com) platform using the original NMR ensemble as input.

As different ΔΔG sign conventions are used for labeling the stabilizing and destabilizing mutations, the sign was adjusted (ΔΔG < 0 stabilizing, ΔΔG > 0 destabilizing) in the present study in order to enable clear comparison. Only the values for the mutants relative to wild-type are of interest, as the absolute values are not very meaningful. A short summary of the used structure-based methods is given in **Table S2**.

### Mutation analysis using sequence-based methods

The ATP7B protein sequence was obtained from Uniprot (ID P35670 ATP7B_HUMAN). The obtained sequence was then used to perform saturated mutagenesis with several state-of-the-art sequence-based disease-prediction methods, EASE-MM,^71^ PolyPhen-2,^33^ SIFT,^72^ Envision,^73^ PROVEAN,^74^ SNAP.2,^75^ and FATHMM^76^, using the corresponding default settings. For all the sequence-based methods, only the final scores were collected and used for analysis. A short summary of the used sequence-based methods is also given in **Table S2**.

### Identifying disease causing mutations using ATP7B protein conservation analysis

The degree of evolutionary conservation of an amino acid in a protein reflects a balance between its natural tendency to mutate and the overall need to retain the structural integrity and function of the macromolecule. Conservation analysis was performed using the ConSurf server designed for estimating the evolutionary conservation of amino/nucleic acid positions in a protein/DNA/RNA molecule based on comparison to homologous sequences.^77^ The HMMER method was used for homology search with an E-value cutoff of 0.0001against the UNIREF-90 protein database.^78^ The homologs were selected for analysis based on the ConSurf server default criteria, and the resulting allowed amino acid variation at each position was used in the mutation pathogenicity analysis. In addition, the residue classification (buried/exposed) was also considered during analysis as solvent exposure dependencies of the classifications may provide insight not available from the total set of mutations, the typical example being disruptive mutations being more hydrophilic inside the protein but more hydrophobic on the protein surface.

### Co-variation analysis using GREMLIN

For analysis of the functional effects of the mutations in ATP7B, we also built a global statistical model based on the ATP7B multiple sequence alignment to estimate the log-likelihood of any given ATP7B variant. The multiple sequence alignment was established using HHblits in HHsuite with an E-value cutoff of 10^−10^ and four iterations against the uniclust30_2018_08 database.^79^ The obtained multiple sequence alignment was further filtered by removing the sequences (rows) that have more than 50% gap, resulting in 767 homologs. The global statistical model was established based on the obtained ATP7B multiple sequence alignment using the GREMLIN (Beta version 2.1) algorithm implemented in Tensorflow kindly provided by Dr. Sergey Ovchinnikov (FAS Center for Systems Biology, Harvard University).^80–82^ GREMLIN enables the production of statistical models based on both coevolution and conservation data of homologs, which reveals residue contacts and can thus be a powerful estimator of structural-function impacts of amino acid mutations. Please note that the pseudo likelihood optimization in Tensorflow was performed with Adam optimizer because the “LBFGS” optimizer in the original Matlab version of GREMLIN is slow in Tensorflow. The difference in log-likelihood between the wild type and mutant variant allows us to explore the variant pathogenicity considering both site conservation and pairwise co-varying positions, which in general has better accuracy than analyzing each site independently. Similar methods have been previously successfully applied to identify the disease causing mutations.^83^

## Results and Discussion

### Structure- and sequence-based estimation of ATP7B variant pathogenicity

As mentioned above, many ATP7B missense mutations have been linked to WD. The missense mutations may directly impair the ATP7B copper transport or the ATP binding or hydrolysis activity by mutation in the respective binding sites, but could also generally destabilize the membrane protein, partly dissociate it from the membrane or disrupt the trafficking of the protein to the membrane. Random missense mutations are more likely to reduce the thermochemical stability and typically impair the free energy of folding by 1 kcal/mol compared to the wild type, because the protein stability represents an optimized system.^53, 84–87^ Thus, loss of protein stability leading to excessive protein degradation and loss of functional copies of ATP7B protein available for copper transporting is a likely mechanism for WD.

To distinguish these different possibilities, we employed a wide range of both structure-based stability estimation methods and sequence-based disease prediction methods to estimate the consequence of mutations in ATP7B. We used five available NMR structures of different ATP7B protein domains, with the partial structure of the important ATP binding domain shown for illustration in **Figure 1A**. To ensure a valid control test data set, we used the method of exhaustive control mutagenesis^88^ by introducing all possible single-site amino acid substitutions into the wild type sequence, and using the distribution of scores as a control set in an analysis of variance (ANOVA), since properties of pathogenic mutations are meaningless by themselves if not compared to a random or non-pathogenic control set. Our exhaustive mutation control dataset for structure-based methods comprised 11,305 different ATP7B mutations after mapping to the original sequence numbering shown in **Figure 1B** (Uniprot P35670).

**Figure 1.**
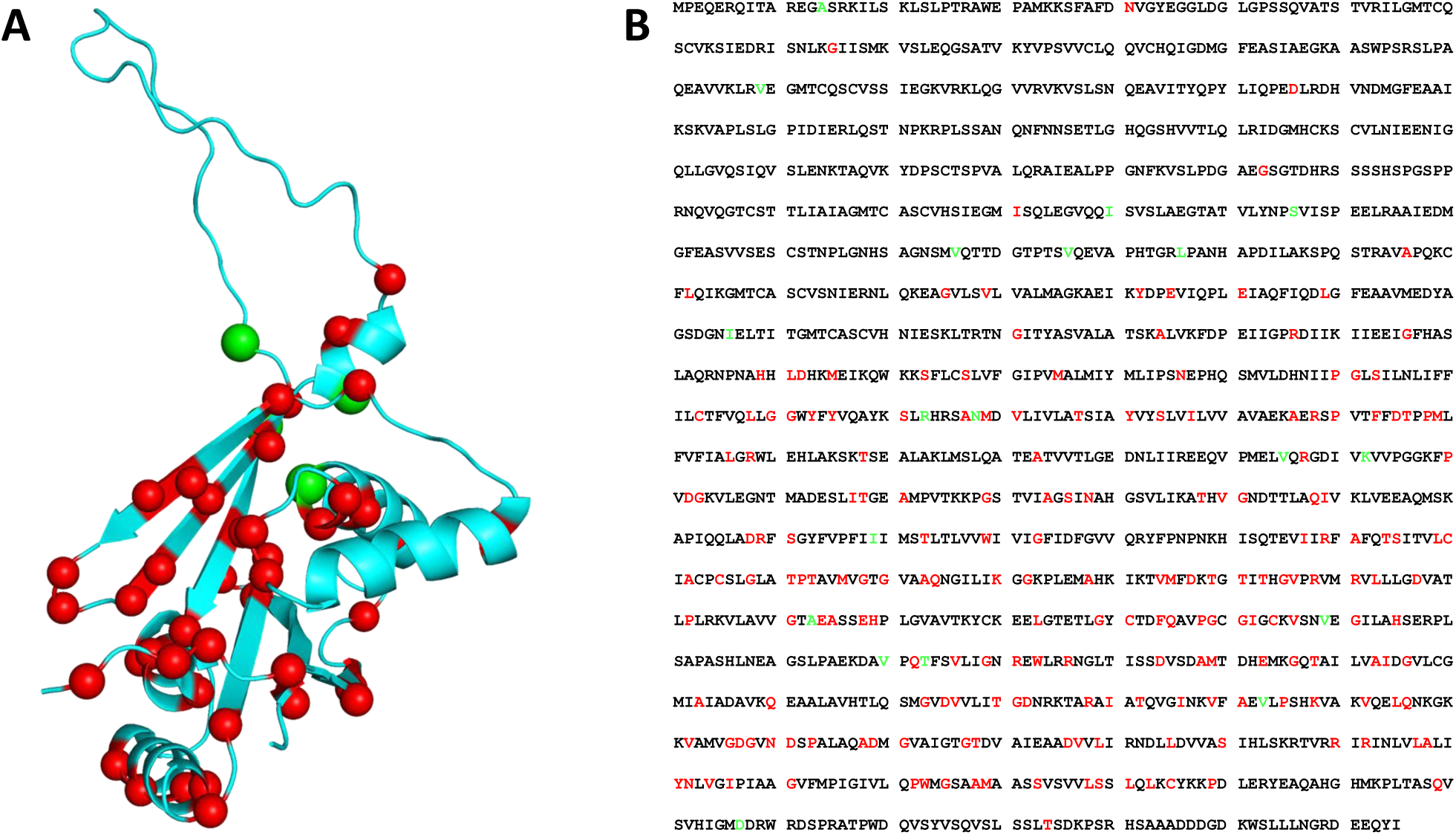
Example structure and full sequence used for in this study. (**A**) The NMR structural model of ATP binding domain (PDB ID 2ARF) as example of used NMR structures of different ATP7B protein domains with pathogenic mutations marked in red and non-pathogenic mutations marked in green. (**B**) The used canonical sequence of ATP7B obtained from Uniprot (ID P35670, ATP7B_HUMAN) with pathogenic mutations marked in red and non pathogenic mutations marked in green.

The computed values for all methods are shown in **Figure 2** with positive ΔΔG values indicating a destabilizing effect on the protein (signs were aligned for all ΔΔG methods to enable comparison). As seen from **Figure 2**, most of the pathogenic mutations destabilize the protein structure. However, the non-pathogenic mutations have similar effects, as expected for randomly introduced mutations,^89^ clearly showing why conclusions cannot be drawn from a set of pathogenic mutations without a proper control. According to the ANOVA summarized in **Table S3**, the destabilization of the non-pathogenic and pathogenic mutations is not significantly different.

**Figure 2.**
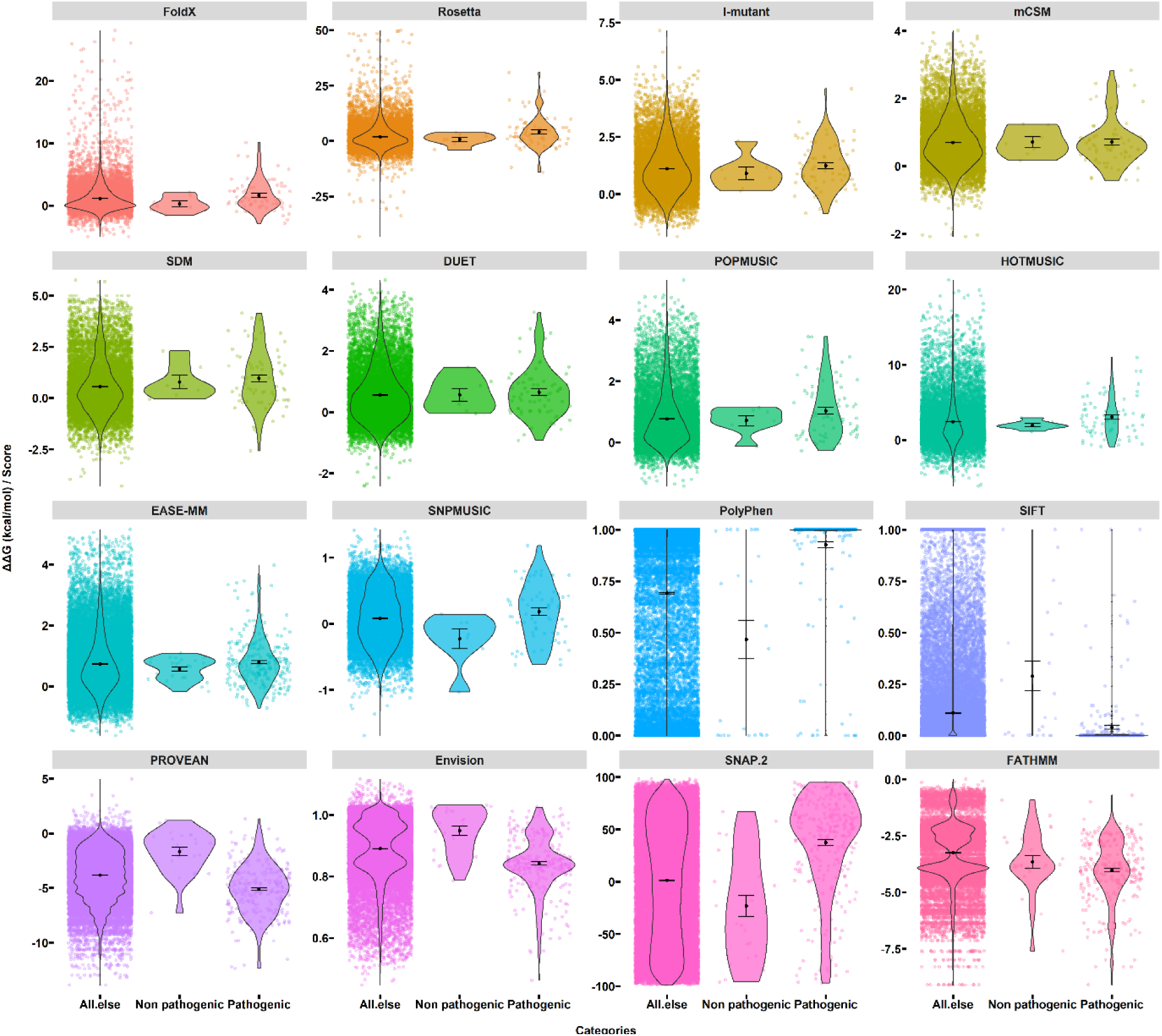
The ΔΔG values/scores for structure-based methods (FoldX, Rosetta, I-mutant, mCSM, SDM, DUET, POPMUSIC, HOTMUSIC, and SNPMUSIC) and sequence-based methods (EASE-MM, Polyphen-2, SIFT, PROVEAN, Envision, SNAP.2, and FATHMM) applied to 291 missense ATP7B mutations. The background dots with jitter function represent the obtained values. The black lines represent the distribution of values. Black dots represent the mean values. The stability methods have signs aligned, whereas the sequence-based methods show pathogenicity relative to their default ranges.

Current protein stability calculators are not expected to be as accurate for membrane proteins as for soluble proteins according to previous benchmarks, since the membrane environment contributes to the stability effect of the mutation.^90^ Furthermore, some of these methods are structure-dependent and the structural input thus affects outcome substantially, with many snapshot structures required to generate an ensemble in agreement with experiment.^52, 54, 91^ Accordingly, we also tested the sequence-based protein stability predictor EASE-MM, which does not rely on structure and thus is not impaired by potential weaknesses in the NMR structures. However, as seen from **Figure 2** and **Table S3**, it also produced insignificant differences in destabilization of pathogenic and control groups. We conclude that pathogenic ATP7B mutations destabilize the protein broadly, but the mutations are not more destabilizing than random mutations in the same protein. This does not rule out that destabilization can contribute to disease as it is a consistent feature of the mutations.

As an alternative to the hypothesis that thermodynamic destabilization of the folded state drives pathogenicity, we also tested whether sequence-based disease predictors using evolutionary conservation information (likelihood of substitution) can better describe pathogenicity. These tools capture important disruptive effects of mutations on the functional folded protein by considering the magnitude of the chemical perturbation and have the advantage of being applicable to all proteins^76, 92^ and several of them describe pathogenicity well in independent benchmarks.^32^ Because such computational methods are error-prone, using one or two of them is not appropriate as the risk of having biased results and conclusions is large if the effect of choice of method on results is not evaluated broadly.

We selected six methods (PolyPhen-2, SIFT, PROVEAN, Envision, SNAP.2 and FATHMM) to study the ATP7B variants. The resulting scores in **Figure 2** for non-pathogenic and pathogenic mutations are well-separated for all six methods except FATHMM. However, such conclusions can be deceptive due to biases in the thresholds, and thus one cannot conclude anything without statistical significance tests. If clinically confirmed non-pathogenic mutations are available at substantial count (> 10) they serve as a relevant test data set in a t-test or ANOVA. Since many data sets do not effectively separate non-pathogenic (benign) variants from confirmed pathogenic variants, we have advocated the use of exhaustive computational mutagenesis as a control data set for t-tests and ANOVA^88^ to test whether a chemical property is significantly different for pathogenic and randomly occurring mutations. Such tests are easily performed using computational methods and provide a statistical quality that is hard to obtain experimentally, due to the cost of functional assaying of thousands of possible mutants. The ANOVA results (**Table S3**) show that the mean values of the obtained scores differ significantly between non-pathogenic and pathogenic categories at the 99% confidence level (p < 0.01) except for the FATHMM method.

In order to test the ability of the methods to distinguish non-pathogenic and pathogenic *ATP7B* mutations, we performed a receiver operating characteristic analysis (**Figure 3A**). In agreement with ANOVA results, the top-5 methods were all sequence-based methods. PolyPhen-2 and PROVEAN produced the highest area under curve (AUC) values of 0.88. Combined with the accuracy results in **Figure 3B**, PROVEAN was slightly (but insignificantly) more accurate than PolyPhen-2. Other properties of the ROC analysis are shown in **Table S4**. The more accurate methods also predict well the pathogenicity of membrane protein mutations, indicating their value when structures are elusive.^32^ One of the most important findings however is that the optimized threshold values for distinguishing pathogenic and non-pathogenic mutations differ widely from the default values of the methods once optimized against the data of ATP7B/WD (**Table 1**). This suggests that studies of disease-related variants need to have the threshold optimized against data for the specific protein of interest (or a class of proteins behaving similarly) as the threshold will be protein-dependent.

**Figure 3.**
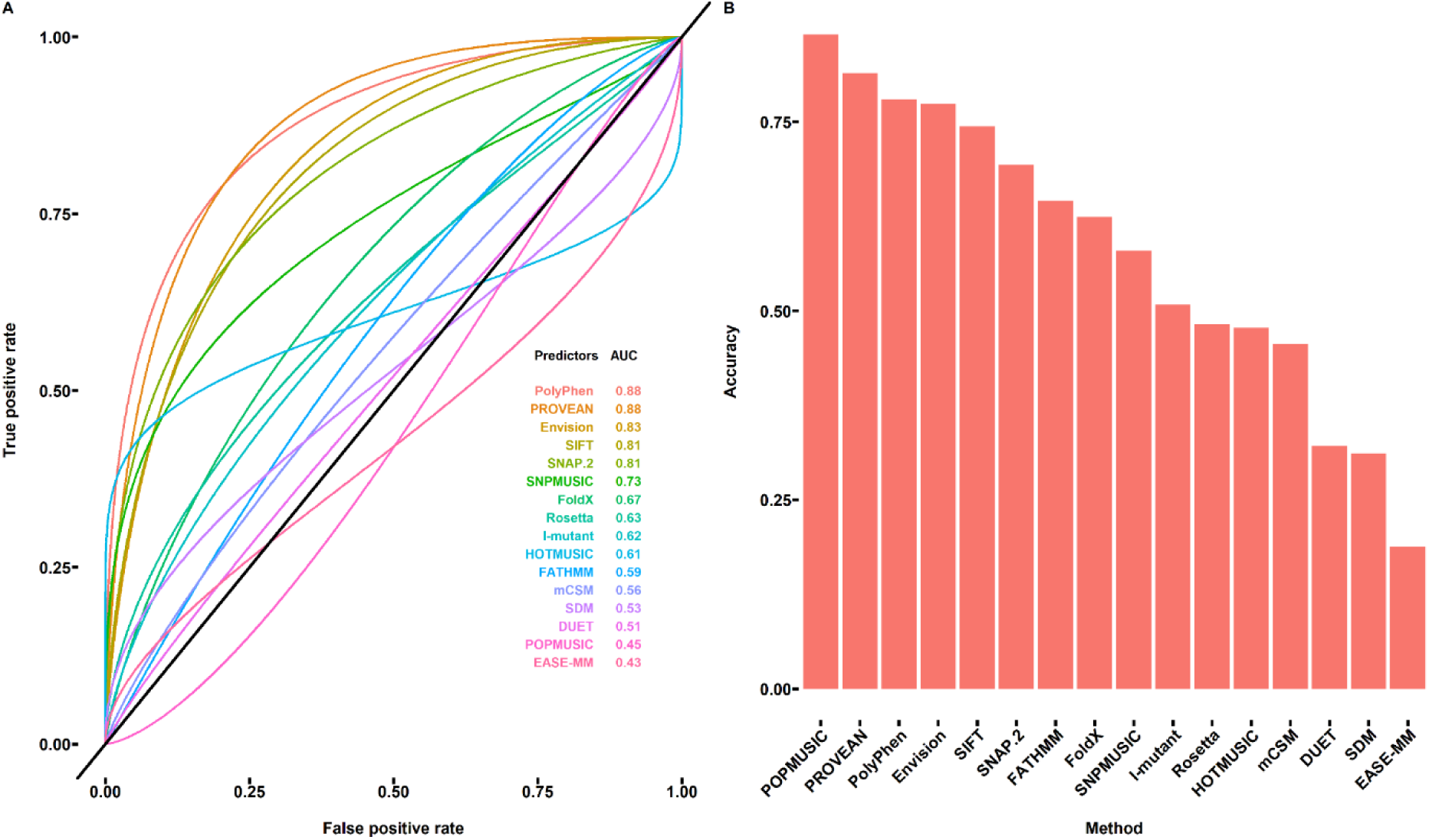
The ROC analysis of methods applied to 291 missense ATP7B mutations. (**A**) The ROC plot of the benchmarked methods for identifying the pathogenicity of the ATP7B protein mutations. (**B**) The identification accuracy of the used methods obtained from ROC analysis.

**Table 1.** Optimized and default threshold for pathogenicity assignment and the number of the pathogenic and non-pathogenic mutations based on the applied threshold.

| Method | Threshold<br>(ROC) | Non<br>Pathogenic | Pathogenic | Threshold<br>(Default) | Non<br>Pathogenic | Pathogenic |
| --- | --- | --- | --- | --- | --- | --- |
| DUET | 1.465 | 9596 | 1705 | 0 | 3238 | 8063 |
| EASE-MM | 0.115 | 6703 | 21132 | 0 | 5093 | 22742 |
| Envision | 0.864 | 16415 | 11420 | 0.5 | 27826 | 9 |
| FATHMM | -3.945 | 19980 | 7855 | -1.5 | 1567 | 26268 |
| FoldX | -0.154 | 2885 | 8420 | 0 | 3570 | 7735 |
| HOTMUSIC | 2.965 | 7869 | 3436 | 0 | 1721 | 9584 |
| I-mutant | 1.180 | 6607 | 4698 | 0 | 1447 | 9858 |
| mCSM | 0.163 | 2553 | 8752 | 0 | 1632 | 9673 |
| PolyPhen | 0.872 | 10827 | 17008 | 0.5 | 8430 | 19405 |
| POPMUSIC | 1.155 | 8444 | 2861 | 0 | 1920 | 9385 |
| PROVEAN | -2.795 | 10151 | 17684 | -2.5 | 8782 | 19053 |
| Rosetta | 3.976 | 8488 | 2790 | 0 | 4368 | 6910 |
| SDM | 0.650 | 6900 | 4401 | 0 | 4034 | 7267 |
| SIFT | 0.0245 | 11312 | 16523 | 0.05 | 9227 | 18608 |
| SNAP.2 | 5.50 | 13757 | 14078 | 0 | 12949 | 14886 |
| SNPMUSIC | 0.185 | 6745 | 4560 | 0 | 4988 | 6317 |

### Correlations between clinical pathogenicity, amino acid properties, and allele frequency

In order to understand how ATP7B mutations confer pathogenicity and to identify valid semi-quantitative prediction tools of ATP7B variant pathogenicity, we analyzed the relationship between the clinically established pathogenicity and changes in 48 amino acid properties previously analyzed in similar work^93^ (summarized in **Table S5**). We calculated the mutation-induced change in property using the equation ΔP = P_mut_ - P_wt_ for both non-pathogenic and pathogenic mutations. Representative results are shown in **Figure 4**, with remaining results shown in **Figure S1**. As illustrated in **Figure 4** and **Figure S1**, most of these properties show no significant separation between non-pathogenic and pathogenic mutations. Based on the ANOVA (**Table S6**) only changes in amino acid bulkiness (B1 in **Figure 4**), the ratio of the side chain volume to its length, differed significantly (95% confidence) for the two categories of mutations, with the pathogenic mutations displaying larger changes in amino acid bulkiness. Hydrophobicity and other changes relating to aggregation propensity, which drive pathogenicity of other disease-causing mutations^88, 94–98^ was not observed for the ATP7B variants, which is in accordance with a view that two types of molecular pathogenicity occurs, one that manifests in soluble proteins subject to aggregation toxicity, and one that manifests in membrane proteins and some soluble allosteric proteins by conformation changes that affect function, distinct from destabilization/aggregation. Previous work suggests that the amino acids bulkiness defines the local conformation and dynamics of natively folded proteins relevant to normal and pathological processes.^99^ Mutation at position 653 of ATP7B indicated that bulky or charged amino acids mimic the phenotype of WD mutations, while small neutral substitutions do not, suggesting that the bulky substitutions distort the ATP7B protein conformation and thereby its function.^100^ Our quantified changes in bulkiness significantly separate pathogenic and non-pathogenic ATP7B mutations for our large data set, suggesting that this hypothesis^100^ is valid for the ATP7B mutations broadly and not just in single cases.

**Figure 4.**
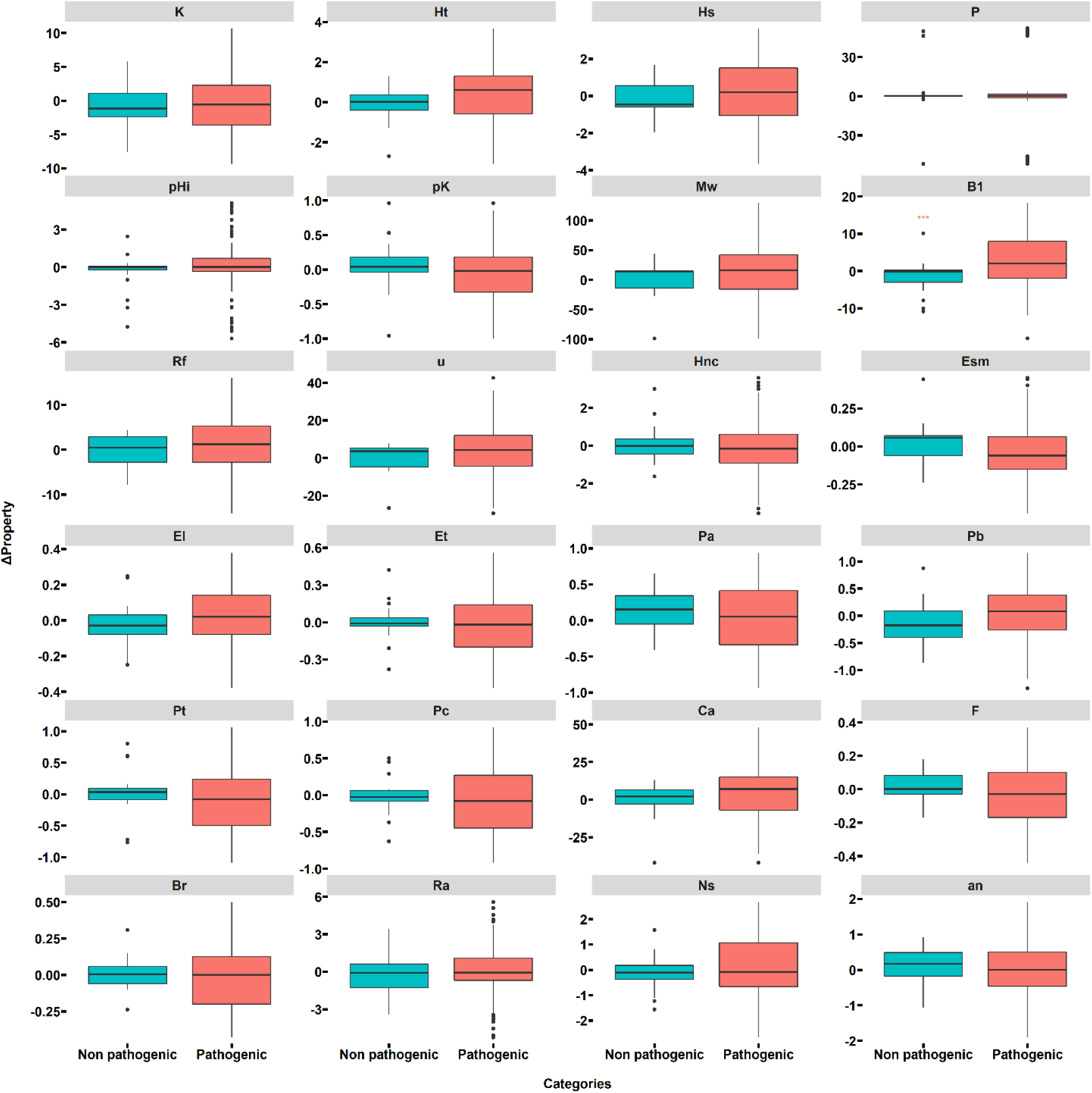
Amino acid property changes for 291 missense ATP7B non-pathogenic and pathogenic ATP7B mutations. Thick bars indicate the medians; the edges of the color-filled rectangles represent the 25^th^ and 75^th^ percentiles. The black dots represent the outliers of the range covered by the black bars. *** for B1 indicates a p-value < 0.05. K: compressibility; Ht: thermodynamic transfer hydrophobicity; Hp: surrounding hydrophobicity; P: polarity; pHi: isoelectric point; pK: equilibrium constant for the ionization of the COOH group; Mw: molecular weight; B1: bulkiness; Rf: chromatographic index; u: refractive index; Hnc: normalized consensus hydrophobicity; Esm: short- and medium-range non-bonded energy; El: long-range non bonded energy; Et: total non bonded energy (Esm + El); Pa, Pb, Pt, and Pc are α-helix, β-strand, turn, and coil tendencies; Ca: helical contact area; F: root-mean-square fluctuational displacement; Br: buriedness; Ra: solvent-accessible reduction ratio; Ns: average number of surrounding residues; an: empirical tendency of the amino acid to be N-terminal.

The database tool gnomAD^101^ estimates the combined allele frequency of ATP7B variants in general populations and thus enables an analysis of the likely natural selection on missense ATP7B mutations across the allele frequency spectrum. The allele frequency is commonly used for clinical diagnostic filtration and is useful for identifying the pathogenicity of some mutations because disease-causing mutations are, all-else being equal, expected to be selected against.

To understand the relationship between pathogenicity and allele frequency for WD mutations, the correlations between screening scores (PolyPhen-2, PROVEAN, Envision, and SNAP.2) and allele frequency for the ATP7B missense mutations found in gnomAD were analyzed. The results in **Figure 5** show that mutations with very high allele frequency are more likely to be benign as correctly predicted by these four methods, in agreement with previous analysis for important Alzheimer’s disease causing mutations in presenilin 1,^32^ which is also a membrane protein with likely metal ion transport function.^102, 103^ Moreover, some non-pathogenic mutations that are misidentified by these methods also had relatively high allele frequency, suggesting that incorporation of this information could improve the prediction of the pathogenicity of ATP7B variants.

**Figure 5.**
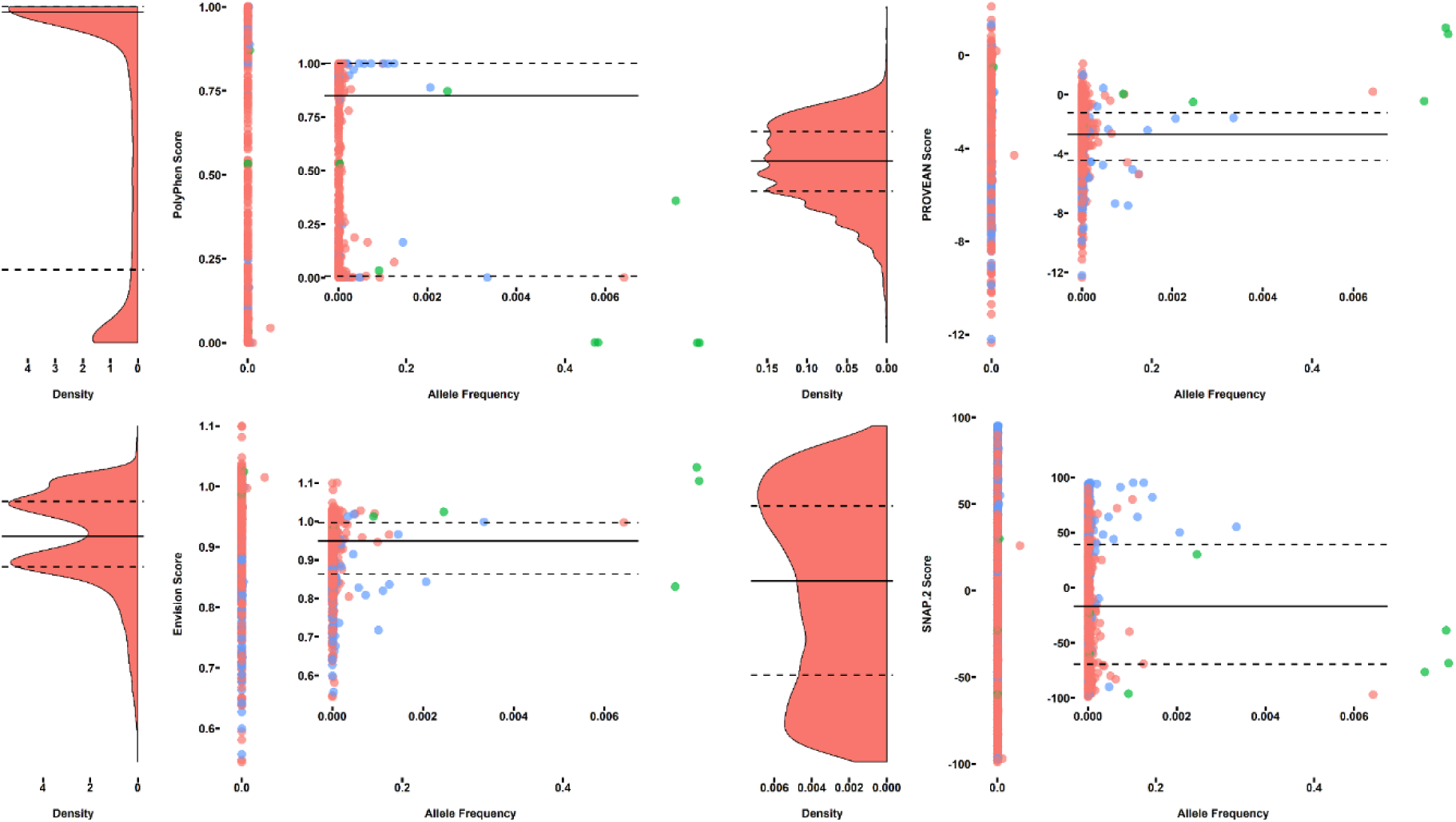
The correlation between PolyPhen, PROVEAN, Envision, and SNAP.2 scores and allele frequency. The plots on the left side in each category are the density plots of the scores. The inset plots on the right side are the zoom-in correlation plots with the allele frequency lower than 0.009. The red, green and blue dots represent the ATP7B protein mutations with unknown pathogenicity, non-pathogenic mutations, and pathogenic mutations, respectively.

### Conservation and co-variation analysis

As we have shown above, several computational methods are capable of predicting the pathogenicity of ATP7B variants at a promising level of accuracy, but only after optimization of their thresholds against a clinically confirmed data set. These sequence-based methods all use evolutionary conservation information. To understand the contribution of this feature in more detail, we analyzed the evolutionary conservation of amino acid positions in ATP7B based on comparison to known ATP7B protein homologs. We extracted the allowed amino acid variations at each position and used this information as a predictor to identify pathogenicity. **Figure 6** shows the results. Simply using the evolutionary conservation information enables good performance with an AUC value of 0.86. This result clearly indicates that conservation information captures an important part of the pathogenic effect on ATP7B protein function.

**Figure 6.**
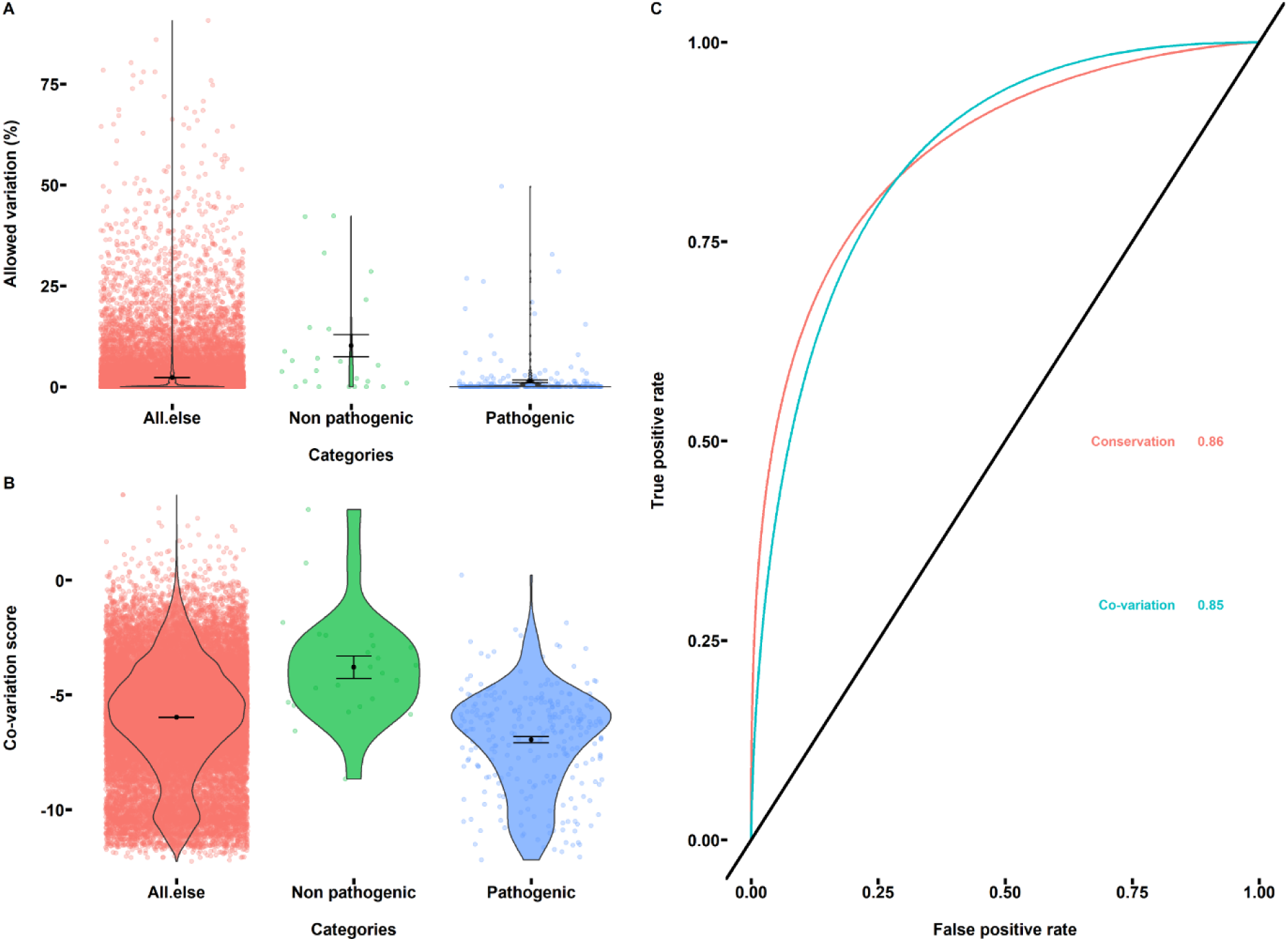
Conservation analysis and co-variation analysis used to identify pathogenic ATP7B mutations. (**A**) The allowed amino acid variation at each position and variation distributions for different categories. The dots in the background represent the obtained allowed variation; the black lines represent the variation distribution. The black dots represent the mean values in each category, and the error bars represent the standard errors in each category. (**B**) The co-variation scores obtained from GREMLIN and score distributions for different categories. (**C**) The ROC plot for conservation analysis and co-variation analysis with the final AUC value labeled.

Previous studies have indicated that most pathogenic mutations occur in buried sites of proteins or in surface-sites involved in molecular interactions.^104–109^ It is thus of interest to know the performance of the computational methods for different types of sites. To explore this, we collected the residue classification information (buried/exposed) based on the evolutionary conservation analysis and divided the ATP7B protein residues into buried and exposed. Then we performed the ROC analysis for all the used methods for each category. As shown in **Figure 7**, it is very clear that most methods perform better for buried residues in agreement with our previous study.^32^ Interestingly, we also found that most non-pathogenic mutations are variable and thus predicted to be exposed, consistent with a neutral effect on protein function.

**Figure 7.**
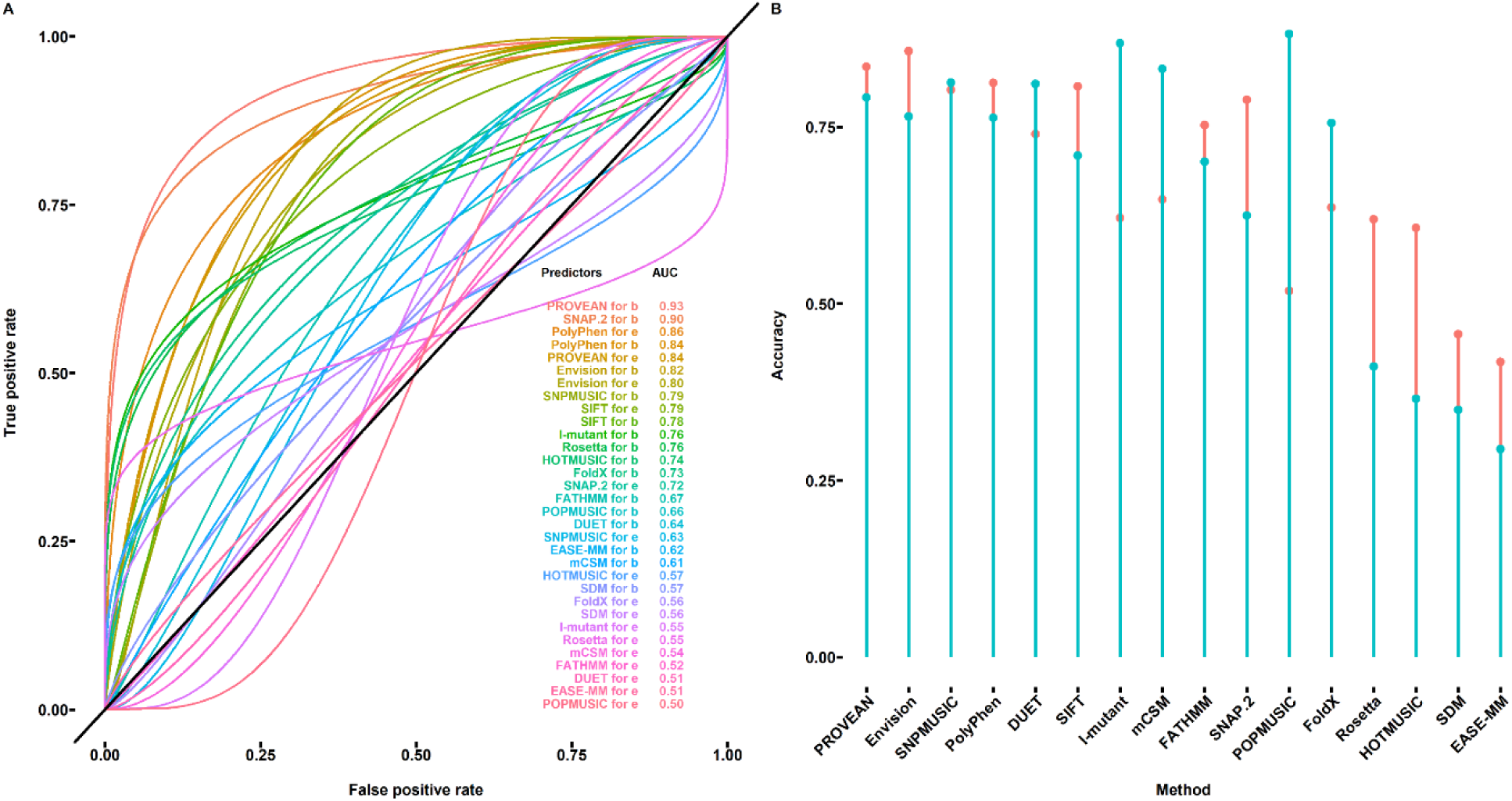
ROC analysis for buried and exposed residues. (**A**) ROC plot of the used structure-based and sequence-based methods applied to predict the pathogenicity of mutations in buried (b) and exposed (e) sites. (**B**) Identification accuracy of the used methods obtained from ROC analysis. The red and green colors represent the accuracy for buried and exposed residues, respectively.

We also analyzed the amino acid property changes for both residue categories using ANOVA (**Table S7**). The difference in the change of amino acid bulkiness for non-pathogenic and pathogenic mutations is only significant for exposed residues, but this may be due to the fact that most of the non-pathogenic mutations are exposed, making the test more assertive for this category, as the significance testing requires a substantial control set. We also find that the β-strand tendency change is significantly different between non-pathogenic and pathogenic mutations but less robust than the amino acids bulkiness. This property has been shown to correlate with protein stability change induced by mutations in some proteins,^93^ and is an ingredient in the aggregation propensity of some proteins.^110^

Since PROVEAN displayed good ability to distinguish the non-pathogenic and pathogenic ATP7B mutations, we investigated this method further. We divided the 20 amino acids into nine groups: Hydrophobic amino acids (AVILM), polar amino acids (SCTQN), aromatic amino acids (FWY), negatively charged (DE), positively charged (HKR), phosphorylatable (STY), small (AGST), proline (P), and glycine (G). Then PROVEAN scores were calculated for each group and residue category (buried/exposed) as shown in **Figure S2**. For buried residues, the mutations related to aromatic amino acids were more likely to be pathogenic. According to the mutation data set shown in **Table S1**, most of the identified pathogenic mutations related to aromatic amino acids substitution were buried and there were no non-pathogenic mutations involving aromatic amino acid substitutions. Based on our analysis above, we think this finding correlates with the bulkiness change identified in the ANOVA analysis. In addition, a previous study indicated that some aromatic amino acids are highly conserved and these positions had to be aromatic amino acids and of a precise size to maintain copper transport function.^111^ For exposed residues, we did not find any systematic tendencies.

So far our analysis has considered each site independently, neglecting the potentially important interactions between residues during substitution. Higher-order statistical models which consider both conservation at individual sites and pairwise coevolution positions may be appropriate to handle this.^31, 81, 112, 113^ These approaches require larger numbers of homologous sequences to build global sequence models.^80, 82^ We applied this method to investigate if such models improve the identification of pathogenic ATP7B variants, as shown in **Figure 6**. We did not see a markedly better performance as indicated by an AUC value of 0.85, suggesting that the interactions between pairs of residues is not critical in driving pathogenicity of ATP7B mutations. When analyzing the conservation and co-variation together, the two methods agree indicating that conservation is the major factor rather than co-variation in the global statistical model.

As shown in **Figure S3**, the ATP binding domain harbors most of the non-pathogenic and pathogenic mutations. The 3D structure of this domain is well defined (PDB ID 2ARF). The relatively small importance of co-variation led us to analyze the structure-based methods again only using the ATP binding domain (**Figure S4**). As seen from the resulting ROC analysis, the AUC value for the best predictor, FoldX, increased to 0.8 indicating that loss of the stability is a relevant driver of disease for the mutations within the ATP binding domain, and thus the failure to identify stability above may be due to poor structural information. The two pathogenic mutations E1064A and H1069Q lower the stability of the ATP binding domain rather than impair ATP binding directly, suggesting a stability rather than functional effect in some of these mutations.^87^

To use our identified drivers in a best-possible combination, we combined the methods to create a two-dimensional representation for the mutations in the ATP binding domain, as shown in **Figure S5**. The individual threshold obtained from ROC analysis is shown as the vertical and horizontal dash lines. It is clear from the analysis that using a two-dimensional representation improves the accuracy of identifying pathogenic mutations, but is still not perfect, and assumptions on the pathogenicity of new variants will clearly be error-prone. Thus, we recommend using the combination of the structure-based approach and conservation analysis to identify the pathogenic ATP7B variants as this two-dimensional representation could greatly improve the accuracy, by e.g. plotting the structure-based output against the sequence-based co-variation data, and optimizing the threshold.

### Sensitivity of findings to data set

Our main finding in this work is that there are major variations in optimal thresholds of the many computational methods, which greatly affect pathogenicity estimates. As argued in the Method section, the WD data is ideal for our purpose as it avoids pollution from computational estimates of the last decade while being nearly complete in confidently assigned mutations (“PS”-type using the Richards et al. classification^38^). To test whether our findings are not affected by the inclusion of recently identified variants, we extracted the missense mutations from the Gao et al. data set from 2019^29^ with likely pathogenicity and performed all the analysis done above also for this data set.

The results are shown in **Figures S6-S12 and Tables S8-S9**. All main findings were unaffected by using the newer data. In particular, our main finding that the computational methods overestimate pathogenicity of ATP7B mutations is unaffected by using the new data, mainly because the confidently assigned missense mutations are similar. In contrast, the total number of reported variants with computationally estimated pathogenicity have increased dramatically the last decade, but were not included to avoid computational self reference in the benchmarking. The sequence-based methods again showed better performance for identifying the pathogenic variants. Furthermore, according to **Figure S8** and **Table S9**, the only significant changes in amino acid properties affecting pathogenicity were again related to amino acid volume or size. For the best-performing sequence-based method PolyPhen and SIFT (those with the highest AUC values), substitutions involving aromatic amino acids tend to confer pathogenicity, consistent with the importance of volume changes inside the membrane protein.

### Implications for estimates of prevalence

The true prevalence of WD has been a longstanding matter of debate: The generally accepted estimate of 1:30,000 rests on a 30-year old publication, whereas recent genetic studies in large populations^15, 29, 30^ have suggested that the true prevalence is in fact four times higher with severe consequences for the large number of undiagnosed patients. These estimates were based on computer evaluations of the variants in the genetic sample. Thus missense mutations were analyzed by Polyphen-2, PhyloP, CADD and MutationTaster in the French study^30^, SIFT and PolyPhen2 in the British study^15^, and SIFT and Polyphen in the global study^29^. Since these authors did not have access to the present evaluation of these methods, we think that they may have overestimated the prevalence by including benign variants. Specifically, the thresholds required to accurately discriminate pathogenic from neutral variants is likely to be very protein-dependent, and unless universal corrections for this fact can be developed, each protein and disease case requires a specific optimization against known clinical data to set the threshold properly in order to estimate disease prevalence. Our study can help to select the best possible model for predicting mutation pathogenicity and thereby yield a more accurate estimate of the genetic prevalence of WD, with the thresholds given in **Table 1**. Using these accurate thresholds may also result in identification of variants with low penetrance, which is crucial for a correct prediction of the number of undiagnosed patients with WD.

Better knowledge of the performance of the computer prediction will also affect the diagnostic work up in patients with suspected WD. According to the Leipzig Criteria,^18^ two disease causing mutations is sufficient for diagnosis which was earlier based on clinical criteria. In such cases it is absolutely important to avoid false diagnoses caused by erroneous computer evaluation of a given variant, a problem that easily arises if thresholds are not optimized as we show here. Considering the likely prevalence of many nearly neutral natural variants, which will probably be clinically benign^38^, we question whether the identification of two mutations is sufficient for the diagnosis without assessment of disturbances in copper metabolism. A similar question arises as concerns the use of genetic testing as mean for neonatal screening for WD, the value of which will heavily depend on the validity of the computer prediction. Our findings regarding the accuracy and thresholds of the applied methods should be important in all of these contexts.

## Conclusions

We have benchmarked state-of-the-art available computational disease prediction methods against a well-known and clinically confirmed data set of pathogenic and non-pathogenic *ATP7B* variants. The data presented suggest that structure-based analysis of the variants does not effectively separate non-pathogenic from pathogenic WD variants whereas sequence-based methods that account for the evolutionary conservation are more effective, but only if their thresholds for distinction are optimized.

Our findings are consistent with a view that proteins (and diseases) differ much more than a default generic threshold a single method can reasonably represent. Thus, different proteins, because of their diversity, have very different thresholds for pathogenicity of an arising mutation, and thus, each method applied should be optimized against a real clinical data set before application. As discussed above, this can affect both diagnosis of WD and the estimation of the real prevalence of the disease. Our results show that prevalence estimates based on these methods are not reliable as they tend to overestimate the pathogenicity of ATP7B mutations. Our finding explains why meta analysis and text search methods^29, 30^, which include many computational estimates, have concluded higher prevalence of WD than clinically observed.

Interestingly, once optimized, the best methods were more effective for buried rather than exposed sites and pointing towards an important role of the bulkiness of the specific amino acid change. The ATP7B protein includes a large transmembrane domain that mediates the transport of copper across the membrane, and buried sites may be those who most likely to affect the transport of copper; our finding that the side-chain volume of buried residues is an important correlator of pathogenicity may be the first structural-functional clue to future more quantitative genotype-phenotype relationships and more accurate prevalence estimates of WD.

## Acknowledgements

The Danish Council for Independent Research | Natural Sciences (DFF), grant case 7016-00079B, and The Memorial Foundation of Manufacturer Vilhelm Petersen & Wife are gratefully acknowledged for supporting this work. The authors are particularly grateful to Dr. Sergey Ovchinnikov for providing the Tensorflow version of GREMLIN and useful suggestions for co-variation analysis.

## Supporting Information

Supporting information is available for this paper and contains details on the analysis including raw data, all p-values from ANOVA, and supplementary figures.

## Supporting Information

**Table S1.** List of non-pathogenic and pathogenic ATP7B missense mutations studied in this work.^1^

| WT | POS | MUT | Area of Protein | Pathology |
| --- | --- | --- | --- | --- |
| N | 41 | S | before Cu1 | Pathogenic |
| G | 85 | V | Cu1 | Pathogenic |
| D | 196 | E | Cu2 | Pathogenic |
| G | 333 | R | bet Cu3/Cu4 | Pathogenic |
| A | 486 | S | Cu5 | Pathogenic |
| L | 492 | S | Cu5 | Pathogenic |
| G | 515 | V | Cu5 | Pathogenic |
| V | 519 | M | Cu5 | Pathogenic |
| Y | 532 | H | Cu5 | Pathogenic |
| V | 536 | A | Cu5 | Pathogenic |
| E | 541 | K | Cu5 | Pathogenic |
| L | 549 | P | Cu5 | Pathogenic |
| G | 591 | S | Cu6 | Pathogenic |
| G | 591 | D | Cu6 | Pathogenic |
| A | 604 | P | Cu6 | Pathogenic |
| R | 616 | W | Cu6 | Pathogenic |
| R | 616 | Q | Cu6 | Pathogenic |
| G | 626 | A | Cu6 | Pathogenic |
| H | 639 | Y | bet Cu6/TM1 | Pathogenic |
| L | 641 | S | bet Cu6/TM1 | Pathogenic |
| D | 642 | H | bet Cu6/TM1 | Pathogenic |
| M | 645 | R | bet Cu6/TM1 | Pathogenic |
| S | 653 | Y | bet Cu6/TM1 | Pathogenic |
| S | 657 | R | TM1 | Pathogenic |
| M | 665 | I | TM1 | Pathogenic |
| N | 676 | I | bet TM1/TM2 | Pathogenic |
| P | 690 | L | TM2 | Pathogenic |
| G | 691 | R | TM2 | Pathogenic |
| S | 693 | P | TM2 | Pathogenic |
| S | 693 | C | TM2 | Pathogenic |
| S | 693 | Y | TM2 | Pathogenic |
| C | 703 | Y | TM2 | Pathogenic |
| L | 708 | P | TM2 | Pathogenic |
| G | 710 | R | TM2 | Pathogenic |
| G | 710 | S | TM2 | Pathogenic |
| G | 710 | A | TM2 | Pathogenic |
| G | 710 | V | TM2 | Pathogenic |
| G | 711 | R | TM2 | Pathogenic |
| G | 711 | W | TM2 | Pathogenic |
| G | 711 | E | TM2 | Pathogenic |
| Y | 713 | C | TM2 | Pathogenic |
| Y | 715 | H | TM2 | Pathogenic |
| S | 721 | P | bet TM2/TM3 | Pathogenic |
| A | 727 | V | bet TM2/TM3 | Pathogenic |
| M | 729 | V | bet TM2/TM3 | Pathogenic |
| V | 731 | E | TM3 | Pathogenic |
| T | 737 | R | TM3 | Pathogenic |
| Y | 741 | C | TM3 | Pathogenic |
| S | 744 | P | TM3 | Pathogenic |
| I | 747 | F | TM3 | Pathogenic |
| A | 756 | G | bet TM3/TM4 | Pathogenic |
| A | 756 | V | bet TM3/TM4 | Pathogenic |
| R | 758 | M | bet TM3/TM4 | Pathogenic |
| P | 760 | L | bet TM3/TM4 | Pathogenic |
| F | 763 | Y | TM4 | Pathogenic |
| D | 765 | G | TM4 | Pathogenic |
| T | 766 | P | TM4 | Pathogenic |
| T | 766 | M | TM4 | Pathogenic |
| T | 766 | R | TM4 | Pathogenic |
| P | 768 | H | TM4 | Pathogenic |
| P | 768 | L | TM4 | Pathogenic |
| M | 769 | R | TM4 | Pathogenic |
| M | 769 | I | TM4 | Pathogenic |
| M | 769 | L | TM4 | Pathogenic |
| L | 776 | P | TM4 | Pathogenic |
| R | 778 | G | TM4 | Pathogenic |
| R | 778 | W | TM4 | Pathogenic |
| R | 778 | Q | TM4 | Pathogenic |
| R | 778 | L | TM4 | Pathogenic |
| L | 795 | F | bet TM4/Td | Pathogenic |
| L | 795 | R | bet TM4/Td | Pathogenic |
| A | 803 | T | bet TM4/Td | Pathogenic |
| R | 827 | P | bet TM4/Td | Pathogenic |
| P | 840 | L | Td | Pathogenic |
| G | 843 | R | Td | Pathogenic |
| D | 842 | N | Td | Pathogenic |
| I | 857 | T | Td | Pathogenic |
| T | 858 | A | Td | Pathogenic |
| A | 861 | T | Td | Pathogenic |
| G | 869 | V | bet Td/TM5 | Pathogenic |
| G | 869 | R | bet Td/TM5 | Pathogenic |
| A | 874 | V | bet Td/TM5 | Pathogenic |
| S | 876 | F | bet Td/TM5 | Pathogenic |
| N | 878 | K | bet Td/TM5 | Pathogenic |
| T | 888 | P | bet Td/TM5 | Pathogenic |
| V | 890 | M | bet Td/TM5 | Pathogenic |
| G | 891 | D | bet Td/TM5 | Pathogenic |
| G | 891 | V | bet Td/TM5 | Pathogenic |
| Q | 898 | R | bet Td/TM5 | Pathogenic |
| I | 899 | F | bet Td/TM5 | Pathogenic |
| D | 918 | N | bet Td/TM5 | Pathogenic |
| D | 918 | A | bet Td/TM5 | Pathogenic |
| D | 918 | E | bet Td/TM5 | Pathogenic |
| R | 919 | G | bet Td/TM5 | Pathogenic |
| R | 919 | W | bet Td/TM5 | Pathogenic |
| S | 921 | N | bet Td/TM5 | Pathogenic |
| T | 933 | P | TM5 | Pathogenic |
| W | 939 | C | TM5 | Pathogenic |
| G | 943 | S | TM5 | Pathogenic |
| G | 943 | C | TM5 | Pathogenic |
| G | 943 | D | TM5 | Pathogenic |
| I | 967 | F | bet TM5/TM6 | Pathogenic |
| R | 969 | W | TM6 | Pathogenic |
| R | 969 | Q | TM6 | Pathogenic |
| A | 971 | V | TM6 | Pathogenic |
| T | 974 | M | TM6 | Pathogenic |
| S | 975 | Y | TM6 | Pathogenic |
| L | 979 | Q | TM6 | Pathogenic |
| C | 980 | Y | TM6 | Pathogenic |
| A | 982 | V | TM6 | Pathogenic |
| C | 985 | Y | TM6 | Pathogenic |
| G | 988 | R | TM6 | Pathogenic |
| T | 991 | M | bet TM6/Ph | Pathogenic |
| P | 992 | L | bet TM6/Ph | Pathogenic |
| P | 992 | H | bet TM6/Ph | Pathogenic |
| T | 993 | M | bet TM6/Ph | Pathogenic |
| M | 996 | T | bet TM6/Ph | Pathogenic |
| G | 998 | D | bet TM6/Ph | Pathogenic |
| G | 1000 | R | bet TM6/Ph | Pathogenic |
| A | 1003 | T | bet TM6/Ph | Pathogenic |
| A | 1003 | V | bet TM6/Ph | Pathogenic |
| Q | 1004 | P | bet TM6/Ph | Pathogenic |
| K | 1010 | R | bet TM6/Ph | Pathogenic |
| K | 1010 | T | bet TM6/Ph | Pathogenic |
| G | 1012 | R | bet TM6/Ph | Pathogenic |
| G | 1012 | V | bet TM6/Ph | Pathogenic |
| A | 1018 | V | bet TM6/Ph | Pathogenic |
| V | 1024 | A | Ph | Pathogenic |
| M | 1025 | R | Ph | Pathogenic |
| D | 1027 | A | Ph | Pathogenic |
| T | 1029 | A | Ph | Pathogenic |
| T | 1029 | I | Ph | Pathogenic |
| T | 1031 | S | Ph | Pathogenic |
| T | 1031 | A | Ph | Pathogenic |
| T | 1033 | A | Ph | Pathogenic |
| T | 1033 | S | Ph | Pathogenic |
| G | 1035 | V | Ph | Pathogenic |
| R | 1038 | K | Ph | Pathogenic |
| R | 1041 | W | ATP loop | Pathogenic |
| R | 1041 | P | ATP loop | Pathogenic |
| L | 1043 | P | ATP loop | Pathogenic |
| D | 1047 | V | ATP loop | Pathogenic |
| P | 1052 | L | ATP loop | Pathogenic |
| G | 1061 | E | ATP loop | Pathogenic |
| E | 1064 | K | ATP loop | Pathogenic |
| E | 1064 | A | ATP loop | Pathogenic |
| A | 1065 | P | ATP loop | Pathogenic |
| E | 1068 | G | ATP loop | Pathogenic |
| H | 1069 | Q | ATP loop | Pathogenic |
| L | 1083 | F | ATP loop | Pathogenic |
| G | 1089 | E | ATP loop | Pathogenic |
| G | 1089 | V | ATP loop | Pathogenic |
| C | 1091 | Y | ATP loop | Pathogenic |
| F | 1094 | L | ATP loop | Pathogenic |
| Q | 1095 | P | ATP loop | Pathogenic |
| P | 1098 | R | ATP loop | Pathogenic |
| G | 1099 | S | ATP loop | Pathogenic |
| G | 1101 | R | ATP loop | Pathogenic |
| I | 1102 | T | ATP loop | Pathogenic |
| C | 1104 | F | ATP loop | Pathogenic |
| C | 1104 | Y | ATP loop | Pathogenic |
| V | 1106 | D | ATP loop | Pathogenic |
| G | 1111 | D | ATP loop | Pathogenic |
| R | 1115 | H | ATP loop | Pathogenic |
| Q | 1142 | H | ATP loop | Pathogenic |
| V | 1146 | M | ATP loop | Pathogenic |
| G | 1149 | A | ATP loop | Pathogenic |
| R | 1151 | C | ATP loop | Pathogenic |
| R | 1151 | H | ATP loop | Pathogenic |
| W | 1153 | R | ATP loop | Pathogenic |
| W | 1153 | C | ATP loop | Pathogenic |
| R | 1156 | H | ATP loop | Pathogenic |
| D | 1164 | N | ATP loop | Pathogenic |
| A | 1168 | S | ATP loop | Pathogenic |
| A | 1168 | P | ATP loop | Pathogenic |
| M | 1169 | T | ATP loop | Pathogenic |
| E | 1173 | K | ATP loop | Pathogenic |
| E | 1173 | G | ATP loop | Pathogenic |
| G | 1176 | R | ATP loop | Pathogenic |
| G | 1176 | E | ATP loop | Pathogenic |
| T | 1178 | A | ATP loop | Pathogenic |
| A | 1183 | T | ATP loop | Pathogenic |
| G | 1186 | S | ATP loop | Pathogenic |
| G | 1186 | C | ATP loop | Pathogenic |
| A | 1193 | P | ATP loop | Pathogenic |
| Q | 1200 | P | ATP loop | Pathogenic |
| G | 1213 | V | ATP bind | Pathogenic |
| D | 1215 | Y | ATP bind | Pathogenic |
| V | 1216 | M | ATP bind | Pathogenic |
| T | 1220 | M | ATP bind | Pathogenic |
| G | 1221 | E | ATP bind | Pathogenic |
| D | 1222 | N | ATP bind | Pathogenic |
| D | 1222 | Y | ATP bind | Pathogenic |
| D | 1222 | V | ATP bind | Pathogenic |
| R | 1228 | T | ATP bind | Pathogenic |
| I | 1230 | V | ATP bind | Pathogenic |
| T | 1232 | P | ATP bind | Pathogenic |
| I | 1236 | T | ATP bind | Pathogenic |
| V | 1239 | G | ATP bind | Pathogenic |
| A | 1241 | V | ATP hinge | Pathogenic |
| K | 1248 | N | ATP hinge | Pathogenic |
| V | 1252 | I | ATP hinge | Pathogenic |
| L | 1255 | I | ATP hinge | Pathogenic |
| Q | 1256 | R | ATP hinge | Pathogenic |
| V | 1262 | F | ATP hinge | Pathogenic |
| G | 1266 | R | ATP hinge | Pathogenic |
| G | 1266 | K | ATP hinge | Pathogenic |
| G | 1266 | V | ATP hinge | Pathogenic |
| D | 1267 | A | ATP hinge | Pathogenic |
| D | 1267 | V | ATP hinge | Pathogenic |
| G | 1268 | R | ATP hinge | Pathogenic |
| N | 1270 | T | ATP hinge | Pathogenic |
| N | 1270 | S | ATP hinge | Pathogenic |
| D | 1271 | N | ATP hinge | Pathogenic |
| P | 1273 | S | ATP hinge | Pathogenic |
| P | 1273 | L | ATP hinge | Pathogenic |
| P | 1273 | Q | ATP hinge | Pathogenic |
| A | 1278 | V | ATP hinge | Pathogenic |
| D | 1279 | Y | ATP hinge | Pathogenic |
| D | 1279 | G | ATP hinge | Pathogenic |
| G | 1281 | C | ATP hinge | Pathogenic |
| G | 1281 | D | ATP hinge | Pathogenic |
| G | 1287 | S | ATP hinge | Pathogenic |
| T | 1288 | R | ATP hinge | Pathogenic |
| T | 1288 | M | ATP hinge | Pathogenic |
| D | 1296 | N | bet ATP hinge/TM7 | Pathogenic |
| V | 1297 | D | bet ATP hinge/TM7 | Pathogenic |
| L | 1299 | F | bet ATP hinge/TM7 | Pathogenic |
| L | 1305 | P | bet ATP hinge/TM7 | Pathogenic |
| S | 1310 | R | bet ATP hinge/TM7 | Pathogenic |
| R | 1320 | S | bet ATP hinge/TM7 | Pathogenic |
| R | 1322 | P | bet ATP hinge/TM7 | Pathogenic |
| L | 1327 | V | TM7 | Pathogenic |
| A | 1328 | T | TM7 | Pathogenic |
| Y | 1331 | S | TM7 | Pathogenic |
| N | 1332 | D | TM7 | Pathogenic |
| N | 1332 | K | TM7 | Pathogenic |
| V | 1334 | D | TM7 | Pathogenic |
| I | 1336 | T | TM7 | Pathogenic |
| G | 1341 | S | TM7 | Pathogenic |
| G | 1341 | R | TM7 | Pathogenic |
| G | 1341 | D | TM7 | Pathogenic |
| G | 1341 | V | TM7 | Pathogenic |
| P | 1352 | S | TM8 | Pathogenic |
| P | 1352 | L | TM8 | Pathogenic |
| P | 1352 | R | TM8 | Pathogenic |
| W | 1353 | R | TM8 | Pathogenic |
| G | 1355 | S | TM8 | Pathogenic |
| A | 1358 | S | TM8 | Pathogenic |
| M | 1359 | I | TM8 | Pathogenic |
| S | 1363 | F | TM8 | Pathogenic |
| L | 1368 | P | TM8 | Pathogenic |
| S | 1369 | L | TM8 | Pathogenic |
| L | 1371 | P | TM8 | Pathogenic |
| L | 1373 | P | TM8 | Pathogenic |
| L | 1373 | R | TM8 | Pathogenic |
| C | 1375 | S | TM8 | Pathogenic |
| P | 1379 | S | after TM8 | Pathogenic |
| Q | 1399 | R | after TM8 | Pathogenic |
| T | 1434 | M | after TM8 | Pathogenic |
| I | 381 | S | Cu4 | Pathogenic |
| A | 727 | D | bet TM2/TM3 | Pathogenic |
| I | 1184 | T | ATP loop | Pathogenic |
| G | 988 | V | TM6 | Pathogenic |
| P | 1245 | T | ATP hinge | Pathogenic |
| V | 1036 | I | ATP loop | Pathogenic |
| T | 788 | I | bet TM4/Td | Pathogenic |
| A | 14 | D | before Cu1 | Non pathogenic |
| V | 149 | L | Cu2 | Non pathogenic |
| I | 390 | V | Cu4 | Non pathogenic |
| S | 406 | A | Cu4 | Non pathogenic |
| V | 446 | L | bet Cu4/Cu5 | Non pathogenic |
| L | 456 | V | bet Cu4/Cu5 | Non pathogenic |
| L | 466 | V | bet Cu4/Cu5 | Non pathogenic |
| S | 565 | N | Cu6 | Non pathogenic |
| G | 723 | R | bet TM2/TM3 | Non pathogenic |
| N | 728 | D | bet TM2/TM3 | Non pathogenic |
| V | 825 | L | bet TM4/Td | Non pathogenic |
| K | 832 | R | bet TM4/Td | Non pathogenic |
| V | 864 | I | Td | Non pathogenic |
| I | 929 | V | TM5 | Non pathogenic |
| K | 952 | R | bet TM5/TM6 | Non pathogenic |
| A | 1063 | V | ATP loop | Non pathogenic |
| V | 1109 | M | ATP loop | Non pathogenic |
| V | 1140 | A | ATP loop | Non pathogenic |
| T | 1143 | N | ATP loop | Non pathogenic |
| P | 1245 | S | ATP hinge | Non pathogenic |
| D | 1271 | E | ATP hinge | Non pathogenic |
| V | 1297 | I | bet ATP hinge/TM7 | Non pathogenic |
| D | 1407 | E | after TM8 | Non pathogenic |
| V | 1243 | M | ATP hinge | Non pathogenic |

**Table S2.** A summary of the used structure-based and sequence-based methods.

| Method | Description | Input | Threshold |
| --- | --- | --- | --- |
| FoldX | ProTherm experimental $\Delta\Delta G$ calibrated empirical force field | 3D structure | $\Delta\Delta G < 0$ Stabilizing<br>$\Delta\Delta G > 0$ Destabilizing |
| Rosetta | Structure knowledge based potential using cartesian space sampling | 3D structure | $\Delta\Delta G < 0$ Stabilizing<br>$\Delta\Delta G > 0$ Destabilizing |
| I-mutant 2.0 | Support vector machine (SVM) based protein stability predictor | 3D structure | $\Delta\Delta G > 0$ Stabilizing<br>$\Delta\Delta G < 0$ Destabilizing |
| mCSM | Graph based structural signatures | 3D structure | $\Delta\Delta G > 0$ Stabilizing<br>$\Delta\Delta G < 0$ Destabilizing |
| SDM | statistical potential energy function | 3D structure | $\Delta\Delta G > 0$ Stabilizing<br>$\Delta\Delta G < 0$ Destabilizing |
| DUET | Support vector machine (SVM) based protein stability predictor combines mCSM and SDM | 3D structure | $\Delta\Delta G > 0$ Stabilizing<br>$\Delta\Delta G < 0$ Destabilizing |
| POPMUSIC | Linear combination of statistical potentials | 3D structure | $\Delta\Delta G < 0$ Stabilizing<br>$\Delta\Delta G > 0$ Destabilizing |
| HOTMUSIC | Thermodynamics based and knowledge driven melting temperature ($\Delta T_m$) predictor | 3D structure | $\Delta T_m > 0$ Stabilizing<br>$\Delta T_m < 0$ Destabilizing |
| EASE-MM | Sequence based prediction with five specialized support vector machine (SVM) models | Sequence | $\Delta\Delta G > 0$ Stabilizing<br>$\Delta\Delta G < 0$ Destabilizing |
| PolyPhen-2 | Naive Bayes classifier trained by HumDiv and HumVar dataset | Sequence | Score $< 0.5$ Benign<br>Score $> 0.5$ Damaging |
| SIFT | Uses sequence homology (PSI-BLAST) combined with known generic likelihood of amino acid substitutions | Sequence | Score $> 0.05$ Tolerated<br>Score $< 0.05$ Deleterious |
| PROVEAN | Uses pairwise sequence alignment scores (BLAST) | Sequence | Score $> -2.5$ Neutral<br>Score $< -2.5$ Deleterious |
| Envision | A predictor of mutational effect magnitude | Sequence | around 1 Neutral<br>around 0 Disruptive |
| SNPMUSIC | Stability driven knowledge based classifier that uses statistical potentials | 3D structure | Score $< 0$ Neutral<br>Score $> 0$ Deleterious |
| SNAP.2 | Neural network (NN) based classifier trained by experimentally obtained variant functional effect data | Sequence | Score $< 0.5$ Neutral<br>Score $> 0.5$ Effect |
| FATHMM | Hidden Markov Models based classifier | Sequence | Score $> -1.5$ Tolerated<br>Score $< -1.5$ Damaging |

**Table S3.**
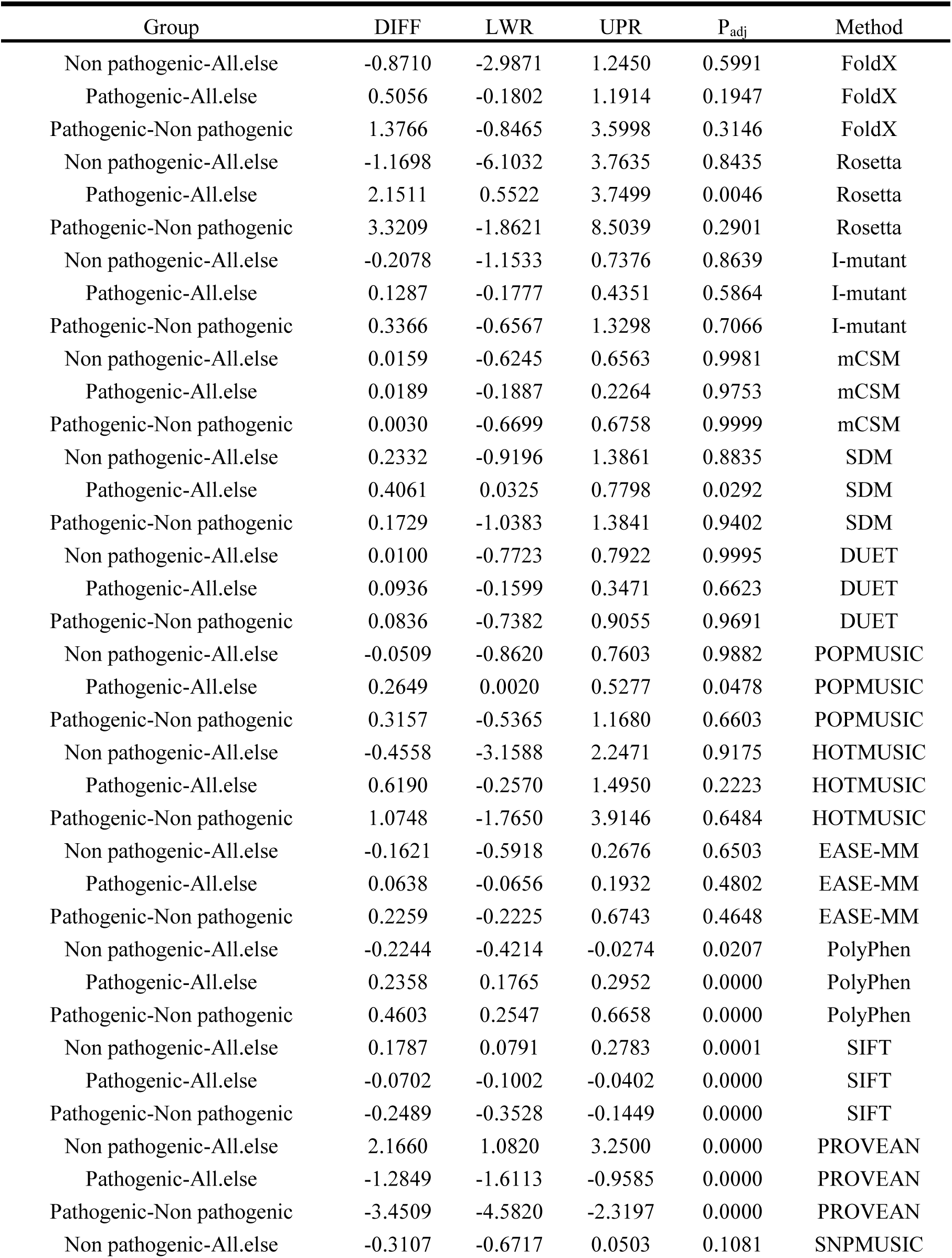
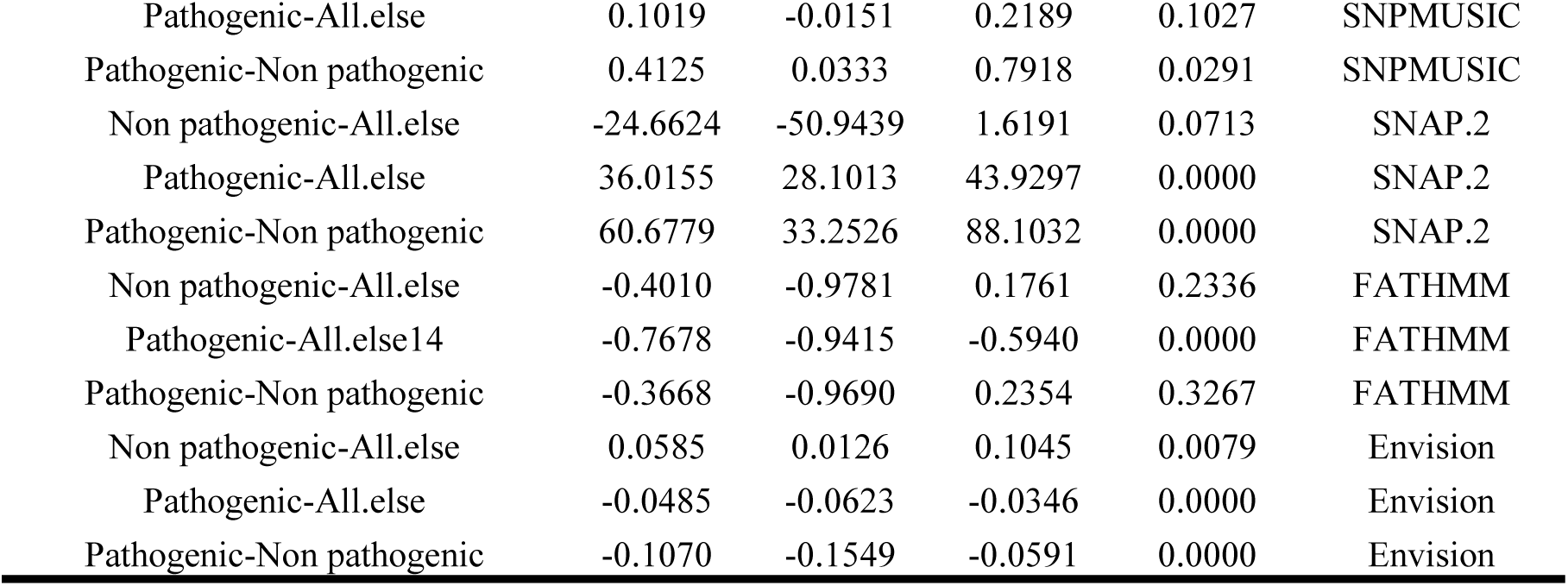
The ANOVA for all the used methods. DIFF refers to the difference between means of the two groups. LWR and UPR represent the lower and the upper end-point of the confidence interval at 95%, respectively. Padj is the adjusted p-value for multiple comparisons. Please note that since different methods use different sign convention for stabilizing and destabilizing, all signs were adjusted to enable comparison for all the protein stability methods such that ΔΔG < 0 is stabilizing, ΔΔG > 0 destabilizing).

**Table S4.** Properties of the ROC analysis.

| Method | specificity | sensitivity | True<br>negative | True<br>positive | False<br>negative | False<br>positive | negative<br>predictive<br>value | Positive<br>predictive<br>value | precision | recall |
| --- | --- | --- | --- | --- | --- | --- | --- | --- | --- | --- |
| PolyPhen | 0.81 | 0.78 | 19.44 | 207.43 | 59.57 | 4.56 | 0.25 | 0.98 | 0.98 | 0.78 |
| PROVEAN | 0.77 | 0.82 | 18.48 | 218.41 | 48.59 | 5.52 | 0.28 | 0.98 | 0.98 | 0.82 |
| Envision | 0.72 | 0.78 | 17.2 | 207.96 | 59.04 | 6.8 | 0.23 | 0.97 | 0.97 | 0.78 |
| SIFT | 0.73 | 0.75 | 17.45 | 199.07 | 67.93 | 6.55 | 0.2 | 0.97 | 0.97 | 0.75 |
| SNAP.2 | 0.78 | 0.68 | 18.76 | 182.88 | 84.12 | 5.24 | 0.18 | 0.97 | 0.97 | 0.68 |
| SNPMUSIC | 0.83 | 0.55 | 5.78 | 37.11 | 29.89 | 1.22 | 0.16 | 0.97 | 0.97 | 0.55 |
| FoldX | 0.62 | 0.62 | 4.34 | 41.83 | 25.17 | 2.66 | 0.15 | 0.94 | 0.94 | 0.62 |
| Rosetta | 0.75 | 0.45 | 5.25 | 30.42 | 36.58 | 1.75 | 0.13 | 0.95 | 0.95 | 0.45 |
| I-mutant | 0.69 | 0.49 | 4.81 | 32.78 | 34.22 | 2.19 | 0.12 | 0.94 | 0.94 | 0.49 |
| HOTMUSIC | 0.94 | 0.43 | 6.61 | 28.71 | 38.29 | 0.39 | 0.15 | 0.99 | 0.99 | 0.43 |
| FATHMM | 0.47 | 0.66 | 11.24 | 176.61 | 90.39 | 12.76 | 0.11 | 0.93 | 0.93 | 0.66 |
| mCSM | 0.65 | 0.44 | 4.52 | 29.24 | 37.76 | 2.48 | 0.11 | 0.92 | 0.92 | 0.44 |
| SDM | 0.87 | 0.25 | 6.12 | 16.91 | 50.09 | 0.88 | 0.11 | 0.95 | 0.95 | 0.25 |
| DUET | 0.75 | 0.28 | 5.27 | 18.49 | 48.51 | 1.73 | 0.1 | 0.91 | 0.91 | 0.28 |
| POPMUSIC | 0.07 | 0.95 | 0.46 | 63.59 | 3.41 | 6.54 | 0.12 | 0.91 | 0.91 | 0.95 |
| EASE-MM | 0.93 | 0.12 | 22.37 | 32.4 | 234.6 | 1.63 | 0.09 | 0.95 | 0.95 | 0.12 |

**Table S5.** Summary of the 48 amino acid properties used in this analysis from previously published work.^2^

| Property | A | D | C | E | F | G | H | I | K | L |
| --- | --- | --- | --- | --- | --- | --- | --- | --- | --- | --- |
| K | -25.5 | -33.12 | -32.82 | -36.17 | -34.54 | -27 | -31.84 | -31.78 | -32.4 | -31.78 |
| Ht | 0.87 | 0.66 | 1.52 | 0.67 | 2.87 | 0.1 | 0.87 | 3.15 | 1.64 | 2.17 |
| Hs | 13.05 | 11.1 | 14.3 | 11.41 | 13.89 | 12.2 | 12.42 | 15.34 | 11.01 | 14.19 |
| P | 0 | 49.7 | 1.48 | 49.9 | 0.35 | 0 | 51.6 | 0.1 | 49.5 | 0.13 |
| pHi | 6 | 2.77 | 5.05 | 5.22 | 5.48 | 5.97 | 7.59 | 6.02 | 9.74 | 5.98 |
| pK | 2.34 | 2.01 | 1.65 | 2.19 | 1.89 | 2.34 | 1.82 | 1.36 | 2.18 | 2.36 |
| Mw | 89 | 133 | 121 | 147 | 165 | 75 | 155 | 131 | 146 | 131 |
| B1 | 11.5 | 11.68 | 13.46 | 13.57 | 19.8 | 3.4 | 13.67 | 21.4 | 15.71 | 21.4 |
| Rf | 9.9 | 2.8 | 2.8 | 3.2 | 18.8 | 5.6 | 8.2 | 17.1 | 3.5 | 17.6 |
| u | 14.34 | 12 | 35.77 | 17.26 | 29.4 | 0 | 21.81 | 19.06 | 21.29 | 18.78 |
| Hnc | 0.62 | 0.9 | 0.29 | -0.74 | 1.19 | 0.48 | -0.4 | 1.38 | -1.5 | 1.06 |
| Esm | 1.4 | 1.16 | 1.37 | 1.16 | 1.14 | 1.36 | 1.22 | 1.19 | 1.07 | 1.32 |
| El | 0.49 | 0.35 | 0.67 | 0.37 | 0.72 | 0.53 | 0.54 | 0.76 | 0.3 | 0.65 |
| Et | 1.9 | 1.52 | 2.04 | 1.54 | 1.86 | 1.9 | 1.76 | 1.95 | 1.37 | 1.97 |
| Pa | 1.42 | 1.01 | 0.7 | 1.51 | 1.13 | 0.57 | 1 | 1.08 | 1.16 | 1.21 |
| Pb | 0.83 | 0.54 | 1.19 | 0.37 | 1.38 | 0.75 | 0.87 | 1.6 | 0.74 | 1.3 |
| Pt | 0.66 | 1.46 | 1.19 | 0.74 | 0.6 | 1.56 | 0.95 | 0.47 | 1.01 | 0.59 |
| Pc | 0.71 | 1.21 | 1.19 | 0.84 | 0.71 | 1.52 | 1.07 | 0.66 | 0.99 | 0.69 |
| Ca | 20 | 26 | 25 | 33 | 46 | 13 | 37 | 39 | 46 | 35 |
| F | 0.96 | 1.14 | 0.87 | 1.07 | 0.69 | 1.16 | 0.8 | 0.76 | 1.14 | 0.79 |
| Br | 0.38 | 0.14 | 0.57 | 0.09 | 0.51 | 0.38 | 0.31 | 0.56 | 0.04 | 0.5 |
| Ra | 3.7 | 2.6 | 3.03 | 3.3 | 6.6 | 3.13 | 3.57 | 7.69 | 1.79 | 5.88 |
| Ns | 6.05 | 4.95 | 7.86 | 5.1 | 6.62 | 6.16 | 5.8 | 7.51 | 4.88 | 7.37 |
| an | 1.59 | 0.53 | 0.33 | 1.45 | 1.14 | 0.53 | 0.89 | 1.22 | 1.13 | 1.91 |
| ac | 1.44 | 2.13 | 0.76 | 2.01 | 1.01 | 0.62 | 0.56 | 0.68 | 0.59 | 0.58 |
| am | 1.22 | 0.56 | 1.53 | 1.28 | 1.13 | 0.4 | 2.23 | 0.77 | 1.65 | 1.05 |
| V0 | 60.46 | 73.83 | 67.7 | 85.88 | 121.48 | 43.25 | 98.79 | 107.72 | 108.5 | 107.75 |
| Nm | 2.11 | 1.8 | 1.88 | 2.09 | 1.98 | 1.53 | 1.98 | 1.77 | 1.96 | 2.19 |
| Nl | 3.92 | 2.85 | 5.55 | 2.72 | 4.53 | 4.31 | 3.77 | 5.58 | 2.79 | 4.59 |
| Hgm | 13.85 | 11.61 | 15.37 | 11.38 | 13.93 | 13.34 | 13.82 | 15.28 | 11.58 | 14.13 |
| ASAD | 104 | 132.2 | 132.5 | 161.9 | 182 | 73.4 | 165.8 | 171.5 | 195.2 | 161.4 |
| ASAN | 33.2 | 62.4 | 17.9 | 81 | 33.1 | 29.2 | 57.7 | 28.3 | 107.5 | 31.1 |
| ΔASA | 70.9 | 69.6 | 114.3 | 80.5 | 148.4 | 44 | 107.9 | 142.7 | 87.5 | 129.8 |
| ΔGh | -0.54 | -2.97 | -1.64 | -3.71 | -1.06 | -0.59 | -3.38 | 0.32 | -2.19 | 0.27 |
| GhD | -0.58 | -6.1 | -1.91 | 7.37 | -1.35 | -0.82 | -5.57 | 0.4 | -5.97 | 0.35 |
| GhN | -0.06 | -3.11 | -0.27 | -3.62 | -0.28 | -0.23 | -2.18 | 0.07 | -1.7 | 0.07 |
| ΔHh | -2.24 | -4.54 | -3.43 | -5.63 | -5.11 | -1.46 | -6.83 | -3.84 | -5.02 | -3.52 |
| TΔSh | 1.7 | 1.57 | 1.79 | 1.92 | 4.05 | 0.87 | 3.45 | 4.16 | 2.83 | 3.79 |
| ΔCh | 14.22 | 2.73 | 9.41 | 3.17 | 39.06 | 4.88 | 20.05 | 41.98 | 17.68 | 38.26 |
| ΔGc | 0.51 | 2.89 | 2.71 | 3.58 | 3.22 | 0.68 | 3.95 | -0.4 | 1.87 | -0.35 |
| ΔHc | 2.77 | 4.72 | 8.64 | 5.69 | 11.93 | 1.23 | 7.64 | 4.03 | 3.57 | 3.69 |
| ΔSc | -2.25 | -1.83 | -5.92 | -2.11 | -8.71 | -0.55 | -3.69 | -4.42 | -1.7 | -4.04 |
| ΔG | -0.02 | -0.08 | 1.08 | -0.13 | 2.16 | 0.09 | 0.56 | -0.08 | -0.32 | -0.08 |
| ΔH | 0.51 | 0.18 | 5.21 | 0.05 | 6.82 | -0.23 | 0.79 | 0.19 | -1.45 | 0.17 |
| TΔS | -0.54 | -0.26 | -4.14 | -0.19 | -4.66 | 0.31 | -0.23 | -0.27 | 1.13 | -0.24 |
| v | 1 | 4 | 2 | 5 | 7 | 0 | 6 | 4 | 5 | 4 |
| s | 0 | 2 | 0 | 3 | 2 | 0 | 2 | 1 | 0 | 2 |
| f | 0 | 2 | 1 | 3 | 2 | 0 | 2 | 2 | 4 | 2 |

**Table S5 explanation:** K: compressibility; Ht: thermodynamic transfer hydrophobicity; Hp: surrounding hydrophobicity; P: polarity; pHi: isoelectric point; pK: equilibrium constant with reference to the ionization property of COOH group; Mw: molecular weight; B1: bulkiness; Rf: chromatographic ...
| Property | M | N | P | Q | R | S | T | V | W | Y |
| --- | --- | --- | --- | --- | --- | --- | --- | --- | --- | --- |
| K | -31.18 | -30.9 | -23.25 | -32.6 | -26.62 | -29.88 | -31.23 | -30.62 | -30.24 | -35.01 |
| Ht | 1.67 | 0.09 | 2.77 | 0 | 0.85 | 0.07 | 0.07 | 1.87 | 3.77 | 2.67 |
| Hs | 13.62 | 11.72 | 11.06 | 11.78 | 12.4 | 11.68 | 12.12 | 14.73 | 13.96 | 13.57 |
| P | 1.43 | 3.38 | 1.58 | 3.53 | 52 | 1.67 | 1.66 | 0.13 | 2.1 | 1.61 |
| pHi | 5.74 | 5.41 | 6.3 | 5.65 | 10.76 | 5.68 | 5.66 | 5.96 | 5.89 | 5.66 |
| pK | 2.28 | 2.02 | 1.99 | 2.17 | 1.81 | 2.21 | 2.1 | 2.32 | 2.38 | 2.2 |
| Mw | 149 | 132 | 115 | 146 | 174 | 105 | 119 | 117 | 204 | 181 |
| B1 | 16.25 | 12.82 | 17.43 | 14.45 | 14.28 | 9.47 | 15.77 | 21.57 | 21.61 | 18.03 |
| Rf | 14.7 | 5.4 | 14.8 | 9 | 4.6 | 6.9 | 9.5 | 14.3 | 17 | 15 |
| u | 21.64 | 13.28 | 10.93 | 17.56 | 26.66 | 6.35 | 11.01 | 13.92 | 42.53 | 31.55 |
| Hnc | 0.64 | -0.78 | 0.12 | -0.85 | -2.53 | -0.18 | -0.05 | 1.08 | 0.81 | 0.26 |
| Esm | 1.3 | 1.18 | 1.24 | 1.12 | 0.92 | 1.3 | 1.25 | 1.25 | 1.03 | 1.03 |
| El | 0.65 | 0.38 | 0.46 | 0.4 | 0.55 | 0.45 | 0.52 | 0.73 | 0.83 | 0.65 |
| Et | 1.96 | 1.56 | 1.7 | 1.52 | 1.48 | 1.75 | 1.77 | 1.98 | 1.87 | 1.69 |
| Pa | 1.45 | 0.67 | 0.57 | 1.11 | 0.98 | 0.77 | 0.83 | 1.06 | 1.08 | 0.69 |
| Pb | 1.05 | 0.89 | 0.55 | 1.1 | 0.93 | 0.75 | 1.19 | 1.7 | 1.37 | 1.47 |
| Pt | 0.6 | 1.56 | 1.52 | 0.98 | 0.95 | 1.43 | 0.96 | 0.5 | 0.96 | 1.14 |
| Pc | 0.59 | 1.37 | 1.61 | 0.87 | 1.07 | 1.34 | 1.08 | 0.63 | 0.76 | 1.07 |
| Ca | 43 | 28 | 22 | 36 | 55 | 20 | 28 | 33 | 61 | 46 |
| F | 0.78 | 1.04 | 1.16 | 1.07 | 1.05 | 1.13 | 0.96 | 0.79 | 0.77 | 1.01 |
| Br | 0.42 | 0.15 | 0.18 | 0.11 | 0.07 | 0.23 | 0.23 | 0.48 | 0.4 | 0.26 |
| Ra | 5.21 | 2.12 | 2.12 | 2.7 | 2.53 | 2.43 | 2.6 | 7.14 | 6.25 | 3.03 |
| Ns | 6.39 | 5.04 | 5.65 | 5.45 | 5.7 | 5.53 | 5.81 | 7.62 | 6.98 | 6.73 |
| an | 1.25 | 0.53 | 0 | 0.98 | 0.67 | 0.7 | 0.75 | 1.42 | 1.33 | 0.58 |
| ac | 0.73 | 0.93 | 2.19 | 1.2 | 0.39 | 0.81 | 1.25 | 0.63 | 1.4 | 0.72 |
| am | 1.47 | 0.93 | 0 | 1.63 | 1.59 | 0.87 | 0.46 | 1.2 | 0.46 | 0.52 |
| V0 | 105.35 | 78.01 | 82.83 | 93.9 | 127.34 | 60.62 | 76.83 | 90.78 | 143.91 | 123.6 |
| Nm | 2.27 | 1.84 | 1.32 | 2.03 | 1.94 | 1.57 | 1.57 | 1.63 | 1.9 | 1.67 |
| Nl | 4.14 | 3.64 | 3.57 | 3.06 | 3.78 | 3.75 | 4.09 | 5.43 | 4.83 | 4.93 |
| Hgm | 13.86 | 13.02 | 12.35 | 12.61 | 13.1 | 13.39 | 12.7 | 14.56 | 15.48 | 13.88 |
| ASAD | 189.8 | 134.9 | 135.1 | 164.9 | 210.2 | 111.4 | 130.4 | 143.9 | 208.8 | 196.4 |
| ASAN | 41.3 | 60.5 | 60.7 | 71.5 | 94.5 | 48.7 | 52 | 28.1 | 39.5 | 50.4 |
| ΔASA | 147.9 | 74 | 73.5 | 93.3 | 116 | 62.8 | 78 | 115.6 | 167.8 | 145.9 |
| ΔGh | -0.6 | -3.55 | 0.32 | -3.92 | -5.96 | -3.82 | -1.97 | 0.13 | -3.8 | -5.64 |
| GhD | -0.71 | -6.63 | 0.56 | -7.12 | -12.78 | -6.18 | -3.66 | 0.18 | -4.71 | -8.45 |
| GhN | -0.1 | -3.03 | 0.23 | -3.15 | -6.85 | -2.36 | -1.69 | 0.04 | -0.88 | -2.82 |
| ΔHh | -4.16 | -5.68 | -1.95 | -6.23 | -10.43 | -5.94 | -4.39 | -3.15 | -8.99 | -10.67 |
| -TΔSh | 3.56 | 2.13 | 2.27 | 2.31 | 4.47 | 2.12 | 2.42 | 3.28 | 5.19 | 5.03 |
| ΔCh | 31.67 | 3.91 | 23.69 | 3.74 | 16.66 | 6.14 | 16.11 | 32.58 | 37.69 | 30.54 |
| ΔGc | 1.13 | 3.26 | -0.39 | 3.69 | 5.25 | 3.42 | 1.74 | -0.19 | 5.59 | 6.56 |
| ΔHc | 7.06 | 3.64 | 1.97 | 4.47 | 6.03 | 5.8 | 4.42 | 3.45 | 13.46 | 14.41 |
| -TΔSc | -5.93 | -0.39 | -2.36 | -0.78 | -0.78 | -2.38 | -2.68 | -3.64 | -7.87 | -7.95 |
| $\Delta G$ | 0.53 | -0.3 | -0.06 | -0.23 | -0.71 | -0.4 | -0.24 | -0.06 | 1.78 | 0.91 |
| $\Delta H$ | 2.89 | -2.03 | 0.02 | -1.76 | -4.4 | -0.16 | 0.04 | 0.3 | 4.47 | 3.73 |
| $-T\Delta S$ | -2.36 | 1.74 | -0.08 | 1.53 | 3.69 | -0.24 | -0.28 | -0.36 | -2.69 | -2.82 |
| v | 4 | 4 | 3 | 5 | 7 | 2 | 3 | 3 | 10 | 8 |
| s | 0 | 2 | 0 | 3 | 5 | 0 | 1 | 1 | 2 | 2 |
| f | 3 | 2 | 0 | 3 | 5 | 1 | 1 | 1 | 2 | 2 |

**Table S6.** The ANOVA for amino acid property changes. DIFF refers to the difference between means of the two groups. LWR and UPR represent the lower and the upper end-point of the confidence interval at 95%, respectively. P_adj_ is the adjusted p-value for multiple comparisons.

| Group | DIFF | LWR | UPR | $P_{adj}$ | Property |
| --- | --- | --- | --- | --- | --- |
| Pathogenic-Non pathogenic | 0.3138 | -1.6174 | 2.2449 | 0.7494 | K |
| Pathogenic-Non pathogenic | 0.4245 | -0.1689 | 1.0179 | 0.1602 | Ht |
| Pathogenic-Non pathogenic | 0.3229 | -0.3633 | 1.0091 | 0.3551 | Hs |
| Pathogenic-Non pathogenic | 1.2072 | -11.1403 | 13.5546 | 0.8475 | P |
| Pathogenic-Non pathogenic | 0.6299 | -0.3949 | 1.6548 | 0.2273 | pHi |
| Pathogenic-Non pathogenic | -0.1018 | -0.2593 | 0.0557 | 0.2042 | pK |
| Pathogenic-Non pathogenic | 13.2172 | -3.7667 | 30.2012 | 0.1267 | Mw |
| Pathogenic-Non pathogenic | 3.7213 | 0.8384 | 6.6042 | 0.0116 | B1 |
| Pathogenic-Non pathogenic | 1.2232 | -1.3575 | 3.8039 | 0.3517 | Rf |
| Pathogenic-Non pathogenic | 3.8801 | -1.0838 | 8.8440 | 0.1250 | u |
| Pathogenic-Non pathogenic | -0.2486 | -0.8378 | 0.3406 | 0.4070 | Hnc |
| Pathogenic-Non pathogenic | -0.0789 | -0.1584 | 0.0007 | 0.0520 | Esm |
| Pathogenic-Non pathogenic | 0.0422 | -0.0226 | 0.1070 | 0.2007 | El |
| Pathogenic-Non pathogenic | -0.0397 | -0.1415 | 0.0621 | 0.4435 | Et |
| Pathogenic-Non pathogenic | -0.0662 | -0.2413 | 0.1089 | 0.4574 | Pa |
| Pathogenic-Non pathogenic | 0.2331 | 0.0272 | 0.4391 | 0.0267 | Pb |
| Pathogenic-Non pathogenic | -0.0880 | -0.3075 | 0.1315 | 0.4308 | Pt |
| Pathogenic-Non pathogenic | -0.0503 | -0.2443 | 0.1437 | 0.6100 | Pc |
| Pathogenic-Non pathogenic | 5.8577 | -1.2065 | 12.9218 | 0.1038 | Ca |
| Pathogenic-Non pathogenic | -0.0445 | -0.1259 | 0.0369 | 0.2831 | F |
| Pathogenic-Non pathogenic | -0.0239 | -0.1113 | 0.0635 | 0.5909 | Br |
| Pathogenic-Non pathogenic | 0.3694 | -0.5751 | 1.3138 | 0.4421 | Ra |
| Pathogenic-Non pathogenic | 0.1888 | -0.2725 | 0.6501 | 0.4212 | Ns |
| Pathogenic-Non pathogenic | -0.1217 | -0.4345 | 0.1912 | 0.4447 | an |
| Pathogenic-Non pathogenic | -0.0443 | -0.3982 | 0.3095 | 0.8055 | ac |
| Pathogenic-Non pathogenic | 0.1100 | -0.2203 | 0.4404 | 0.5127 | am |
| Pathogenic-Non pathogenic | 12.7474 | -1.7603 | 27.2551 | 0.0848 | V0 |
| Pathogenic-Non pathogenic | -0.1276 | -0.3047 | 0.0496 | 0.1575 | Nm |
| Pathogenic-Non pathogenic | 0.3099 | -0.1123 | 0.7321 | 0.1497 | NI |
| Pathogenic-Non pathogenic | 0.2680 | -0.2914 | 0.8273 | 0.3465 | Hgm |
| Pathogenic-Non pathogenic | 17.9738 | -4.2116 | 40.1592 | 0.1119 | ASAD |
| Pathogenic-Non pathogenic | 5.4339 | -7.0736 | 17.9414 | 0.3932 | ASAN |
| Pathogenic-Non pathogenic | 12.5847 | -5.0118 | 30.1812 | 0.1603 | ΔASA |
| Pathogenic-Non pathogenic | -0.4534 | -1.5983 | 0.6916 | 0.4364 | ΔGh |
| Pathogenic-Non pathogenic | -2.0660 | -4.5996 | 0.4676 | 0.1096 | GhD |
| Pathogenic-Non pathogenic | -0.4692 | -1.7454 | 0.8071 | 0.4699 | GhN |
| Pathogenic-Non pathogenic | -1.1214 | -2.6855 | 0.4428 | 0.1593 | ΔHh |
| Pathogenic-Non pathogenic | 0.6680 | -0.0024 | 1.3384 | 0.0508 | -TΔSh |
| Pathogenic-Non pathogenic | 4.9547 | -1.4481 | 11.3574 | 0.1288 | ΔCh |
| Pathogenic-Non pathogenic | 0.4413 | -0.6142 | 1.4967 | 0.4113 | ΔGc |
| Pathogenic-Non pathogenic | 0.8913 | -0.5883 | 2.3710 | 0.2367 | ΔHc |
| Pathogenic-Non pathogenic | -0.4538 | -1.4910 | 0.5834 | 0.3899 | -TΔSc |
| Pathogenic-Non pathogenic | -0.0128 | -0.3339 | 0.3082 | 0.9374 | $\Delta G$ |
| Pathogenic-Non pathogenic | -0.2279 | -1.4926 | 1.0368 | 0.7231 | $\Delta H$ |
| Pathogenic-Non pathogenic | 0.2172 | -0.7621 | 1.1965 | 0.6628 | $-T\Delta S$ |
| Pathogenic-Non pathogenic | 1.0861 | -0.1741 | 2.3464 | 0.0909 | $v$ |
| Pathogenic-Non pathogenic | 0.3713 | -0.5830 | 1.3255 | 0.4445 | $s$ |
| Pathogenic-Non pathogenic | 0.2069 | -0.6677 | 1.0816 | 0.6418 | $f$ |

**Table S7.** The ANOVA for amino acid property changes of buried and exposed residues. DIFF refers to the difference between means of the two groups. LWR and UPR represent the lower and the upper end-point of the confidence interval at 95%, respectively. P_adj_ is the adjusted p-value for multiple comparisons.

| Group | DIFF | LWR | UPR | $P_{adj}$ | Method | Category |
| --- | --- | --- | --- | --- | --- | --- |
| Pathogenic-Non pathogenic | 0.8782 | -0.1071 | 1.8634 | 0.0802 | Ra | Exposed |
| Pathogenic-Non pathogenic | 13.5992 | -3.7877 | 30.9860 | 0.1243 | V0 | Exposed |
| Pathogenic-Non pathogenic | 5.5221 | -3.3010 | 14.3452 | 0.2181 | Ca | Exposed |
| Pathogenic-Non pathogenic | 0.4471 | -0.9954 | 1.8895 | 0.5411 | pHi | Exposed |
| Pathogenic-Non pathogenic | 0.7548 | 0.1429 | 1.3666 | 0.0160 | Hgm | Exposed |
| Pathogenic-Non pathogenic | 16.9385 | -10.6348 | 44.5117 | 0.2266 | ASAD | Exposed |
| Pathogenic-Non pathogenic | -0.0629 | -0.2829 | 0.1571 | 0.5728 | Pa | Exposed |
| Pathogenic-Non pathogenic | -5.1136 | -20.6618 | 10.4347 | 0.5167 | ASAN | Exposed |
| Pathogenic-Non pathogenic | 0.8075 | 0.0027 | 1.6122 | 0.0492 | TΔSh | Exposed |
| Pathogenic-Non pathogenic | 0.3258 | 0.0920 | 0.5596 | 0.0066 | Pb | Exposed |
| Pathogenic-Non pathogenic | 0.1218 | -0.3104 | 0.5540 | 0.5784 | am | Exposed |
| Pathogenic-Non pathogenic | 10.7865 | -10.0551 | 31.6282 | 0.3080 | Mw | Exposed |
| Pathogenic-Non pathogenic | 0.5817 | 0.1265 | 1.0369 | 0.0126 | Nl | Exposed |
| Pathogenic-Non pathogenic | -0.0231 | -1.1899 | 1.1438 | 0.9689 | f | Exposed |
| Pathogenic-Non pathogenic | -0.4990 | -2.5400 | 1.5420 | 0.6297 | ΔHh | Exposed |
| Pathogenic-Non pathogenic | -10.1742 | -26.6273 | 6.2789 | 0.2236 | P | Exposed |
| Pathogenic-Non pathogenic | -0.0026 | -0.1718 | 0.1667 | 0.9761 | pK | Exposed |
| Pathogenic-Non pathogenic | 4.3601 | -1.5064 | 10.2266 | 0.1440 | u | Exposed |
| Pathogenic-Non pathogenic | -0.0942 | -0.3345 | 0.1461 | 0.4396 | Pc | Exposed |
| Pathogenic-Non pathogenic | 0.0565 | -0.0414 | 0.1543 | 0.2559 | Br | Exposed |
| Pathogenic-Non pathogenic | 4.3706 | 1.0967 | 7.6445 | 0.0092 | B1 | Exposed |
| Pathogenic-Non pathogenic | -0.1577 | -1.4785 | 1.1631 | 0.8138 | ΔGc | Exposed |
| Pathogenic-Non pathogenic | -0.1030 | -0.1956 | -0.0105 | 0.0293 | F | Exposed |
| Pathogenic-Non pathogenic | 0.6342 | -0.0734 | 1.3418 | 0.0786 | Ht | Exposed |
| Pathogenic-Non pathogenic | 3.4593 | 0.6245 | 6.2941 | 0.0171 | Rf | Exposed |
| Pathogenic-Non pathogenic | 21.9564 | 2.0892 | 41.8237 | 0.0305 | ΔASA | Exposed |
| Pathogenic-Non pathogenic | 0.4189 | -2.1527 | 2.9906 | 0.7479 | K | Exposed |
| Pathogenic-Non pathogenic | -0.0613 | -0.2740 | 0.1515 | 0.5701 | Nm | Exposed |
| Pathogenic-Non pathogenic | 9.5099 | 2.6171 | 16.4027 | 0.0072 | ΔCh | Exposed |
| Pathogenic-Non pathogenic | -0.1166 | -0.3911 | 0.1580 | 0.4028 | Pt | Exposed |
| Pathogenic-Non pathogenic | 0.9600 | -0.8027 | 2.7227 | 0.2835 | ΔHc | Exposed |
| Pathogenic-Non pathogenic | 0.0937 | -0.6959 | 0.8832 | 0.8150 | Hnc | Exposed |
| Pathogenic-Non pathogenic | -0.2526 | -0.7147 | 0.2095 | 0.2818 | ac | Exposed |
| Pathogenic-Non pathogenic | 0.0075 | -0.3680 | 0.3829 | 0.9687 | an | Exposed |
| Pathogenic-Non pathogenic | -0.0234 | -0.1232 | 0.0765 | 0.6443 | Esm | Exposed |
| Pathogenic-Non pathogenic | 0.7677 | 0.0333 | 1.5020 | 0.0406 | Hs | Exposed |
| Pathogenic-Non pathogenic | 0.0896 | 0.0162 | 0.1630 | 0.0171 | El | Exposed |
| Pathogenic-Non pathogenic | -1.2604 | -4.7159 | 2.1951 | 0.4721 | GhD | Exposed |
| Pathogenic-Non pathogenic | -0.1442 | -1.4309 | 1.1424 | 0.8250 | s | Exposed |
| Pathogenic-Non pathogenic | 0.5126 | -1.1903 | 2.2155 | 0.5528 | GhN | Exposed |
| Pathogenic-Non pathogenic | 0.0624 | -0.0593 | 0.1840 | 0.3127 | Et | Exposed |
| Pathogenic-Non pathogenic | 0.3085 | -1.1556 | 1.7727 | 0.6777 | ΔGh | Exposed |
| Pathogenic-Non pathogenic | 0.7971 | -0.7095 | 2.3037 | 0.2974 | v | Exposed |
| Pathogenic-Non pathogenic | -0.3093 | -1.5505 | 0.9318 | 0.6230 | TΔS | Exposed |
| Pathogenic-Non pathogenic | 0.5344 | 0.0633 | 1.0055 | 0.0265 | Ns | Exposed |
| Pathogenic-Non pathogenic | 0.4631 | -1.1095 | 2.0357 | 0.5614 | ΔH | Exposed |
| Pathogenic-Non pathogenic | -1.1215 | -2.3402 | 0.0971 | 0.0710 | TΔSc | Exposed |
| Pathogenic-Non pathogenic | 0.1496 | -0.2227 | 0.5219 | 0.4283 | ΔG | Exposed |
| Pathogenic-Non pathogenic | 0.0097 | -1.7491 | 1.7685 | 0.9913 | Ra | Buried |
| Pathogenic-Non pathogenic | 6.7118 | -18.9053 | 32.3290 | 0.6053 | V0 | Buried |
| Pathogenic-Non pathogenic | 3.9644 | -7.9671 | 15.8959 | 0.5124 | Ca | Buried |
| Pathogenic-Non pathogenic | 0.6957 | -0.7914 | 2.1828 | 0.3567 | pHi | Buried |
| Pathogenic-Non pathogenic | -0.1406 | -1.0799 | 0.7988 | 0.7678 | Hgm | Buried |
| Pathogenic-Non pathogenic | 10.7446 | -26.6381 | 48.1273 | 0.5708 | ASAD | Buried |
| Pathogenic-Non pathogenic | -0.0597 | -0.3598 | 0.2404 | 0.6948 | Pa | Buried |
| Pathogenic-Non pathogenic | 13.8286 | -4.5022 | 32.1593 | 0.1381 | ASAN | Buried |
| Pathogenic-Non pathogenic | 0.3340 | -0.8621 | 1.5301 | 0.5818 | TΔSh | Buried |
| Pathogenic-Non pathogenic | 0.1358 | -0.2407 | 0.5123 | 0.4771 | Pb | Buried |
| Pathogenic-Non pathogenic | 0.0767 | -0.4649 | 0.6183 | 0.7799 | am | Buried |
| Pathogenic-Non pathogenic | 9.4361 | -19.2125 | 38.0848 | 0.5160 | Mw | Buried |
| Pathogenic-Non pathogenic | 0.0961 | -0.6619 | 0.8542 | 0.8024 | Nl | Buried |
| Pathogenic-Non pathogenic | 0.1624 | -1.1555 | 1.4803 | 0.8079 | f | Buried |
| Pathogenic-Non pathogenic | -1.5017 | -3.9502 | 0.9468 | 0.2274 | ΔHh | Buried |
| Pathogenic-Non pathogenic | 12.8210 | -4.1613 | 29.8032 | 0.1378 | P | Buried |
| Pathogenic-Non pathogenic | -0.2347 | -0.5332 | 0.0637 | 0.1222 | pK | Buried |
| Pathogenic-Non pathogenic | 2.3344 | -6.6394 | 11.3082 | 0.6079 | u | Buried |
| Pathogenic-Non pathogenic | -0.0148 | -0.3489 | 0.3192 | 0.9302 | Pc | Buried |
| Pathogenic-Non pathogenic | -0.0940 | -0.2369 | 0.0489 | 0.1954 | Br | Buried |
| Pathogenic-Non pathogenic | 2.4654 | -2.9146 | 7.8455 | 0.3666 | B1 | Buried |
| Pathogenic-Non pathogenic | 0.9673 | -0.7442 | 2.6789 | 0.2658 | ΔGc | Buried |
| Pathogenic-Non pathogenic | 0.0260 | -0.1198 | 0.1718 | 0.7251 | F | Buried |
| Pathogenic-Non pathogenic | 0.1920 | -0.8623 | 1.2463 | 0.7193 | Ht | Buried |
| Pathogenic-Non pathogenic | -1.5929 | -6.2776 | 3.0919 | 0.5026 | Rf | Buried |
| Pathogenic-Non pathogenic | -2.8403 | -35.6084 | 29.9278 | 0.8642 | ΔASA | Buried |
| Pathogenic-Non pathogenic | 0.2223 | -2.8732 | 3.3178 | 0.8873 | K | Buried |
| Pathogenic-Non pathogenic | -0.2292 | -0.5445 | 0.0861 | 0.1529 | Nm | Buried |
| Pathogenic-Non pathogenic | -1.1917 | -13.2524 | 10.8690 | 0.8454 | ΔCh | Buried |
| Pathogenic-Non pathogenic | -0.0784 | -0.4542 | 0.2974 | 0.6806 | Pt | Buried |
| Pathogenic-Non pathogenic | 0.5391 | -2.1166 | 3.1947 | 0.6888 | ΔHc | Buried |
| Pathogenic-Non pathogenic | -0.4989 | -1.3482 | 0.3503 | 0.2474 | Hnc | Buried |
| Pathogenic-Non pathogenic | 0.2195 | -0.3446 | 0.7835 | 0.4431 | ac | Buried |
| Pathogenic-Non pathogenic | -0.2157 | -0.7553 | 0.3239 | 0.4307 | an | Buried |
| Pathogenic-Non pathogenic | -0.1260 | -0.2496 | -0.0025 | 0.0456 | Esm | Buried |
| Pathogenic-Non pathogenic | -0.0499 | -1.2953 | 1.1956 | 0.9370 | Hs | Buried |
| Pathogenic-Non pathogenic | -0.0084 | -0.1228 | 0.1060 | 0.8843 | El | Buried |
| Pathogenic-Non pathogenic | -2.1367 | -5.7988 | 1.5254 | 0.2507 | GhD | Buried |
| Pathogenic-Non pathogenic | 0.7728 | -0.6184 | 2.1641 | 0.2740 | s | Buried |
| Pathogenic-Non pathogenic | -1.3620 | -3.1557 | 0.4318 | 0.1356 | GhN | Buried |
| Pathogenic-Non pathogenic | -0.1365 | -0.2912 | 0.0181 | 0.0831 | Et | Buried |
| Pathogenic-Non pathogenic | -1.1678 | -2.9491 | 0.6136 | 0.1971 | ΔGh | Buried |
| Pathogenic-Non pathogenic | 1.0319 | -1.1531 | 3.2170 | 0.3521 | v | Buried |
| Pathogenic-Non pathogenic | 0.7604 | -0.8381 | 2.3590 | 0.3486 | TΔS | Buried |
| Pathogenic-Non pathogenic | -0.1321 | -0.9858 | 0.7216 | 0.7601 | Ns | Buried |
| Pathogenic-Non pathogenic | -0.9622 | -3.0731 | 1.1487 | 0.3691 | ΔH | Buried |
| Pathogenic-Non pathogenic | 0.4261 | -1.4146 | 2.2668 | 0.6479 | TΔSc | Buried |
| Pathogenic-Non pathogenic | -0.1998 | -0.7769 | 0.3773 | 0.4949 | ΔG | Buried |

**Figure S1.**
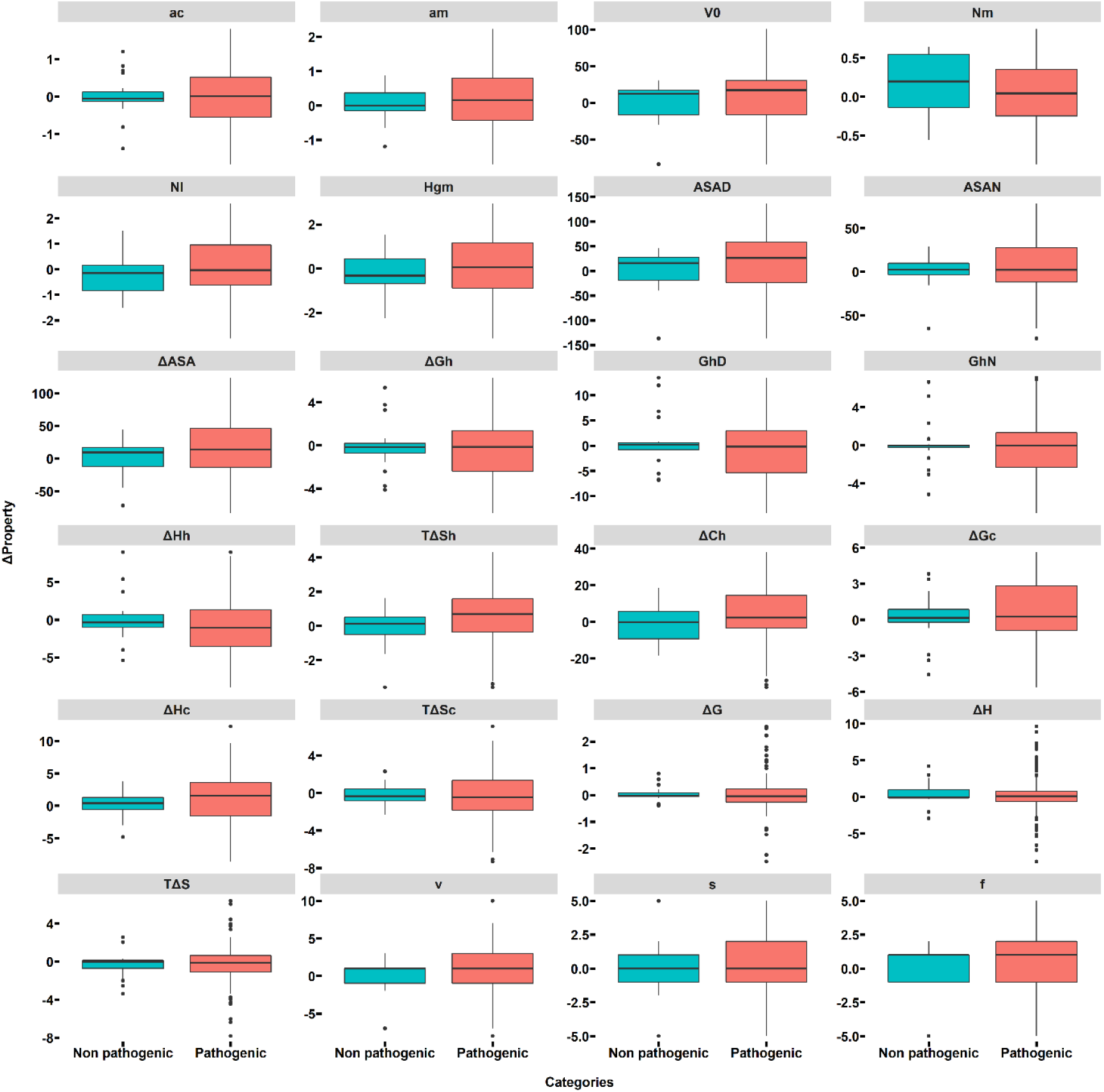
Distribution of amino acid property changes for ATP7B mutations. Thick bars indicate the median, the edges for the color-filled rectangles represent the 25^th^ and 75^th^ percentiles, and the black dots represent the outliers of the range covered by the black bars. ac / am: power to be at the C-terminal and middle of α-helix. V0: partial specific volume; Nm and Nl: average medium- and long-range contacts; Hgm: combined hydrophobicity (globular and membrane); ASAD, ASAN, and ΔASA are solvent-accessible surface area for denatured, native, and unfolding states; ΔGh, ΔGhD, and ΔGhN are the Gibbs free energy change of hydration for unfolding, denatured, and native protein; ΔHh: enthalpy change of hydration; -TΔSh: entropy change of hydration; ΔCh: hydration heat capacity change; ΔGc, ΔHc, and TΔSc are chain-unfolding Gibbs free energy, unfolding enthalpy, and unfolding entropy; ΔG, ΔH, and -TΔS are unfolding Gibbs free energy, enthalpy, and entropy change; v = volume (number of non-hydrogen sidechain atoms); s = shape (position of branch point in a side chain); f = flexibility (number of side-chain dihedral angles).

**Figure S2.**
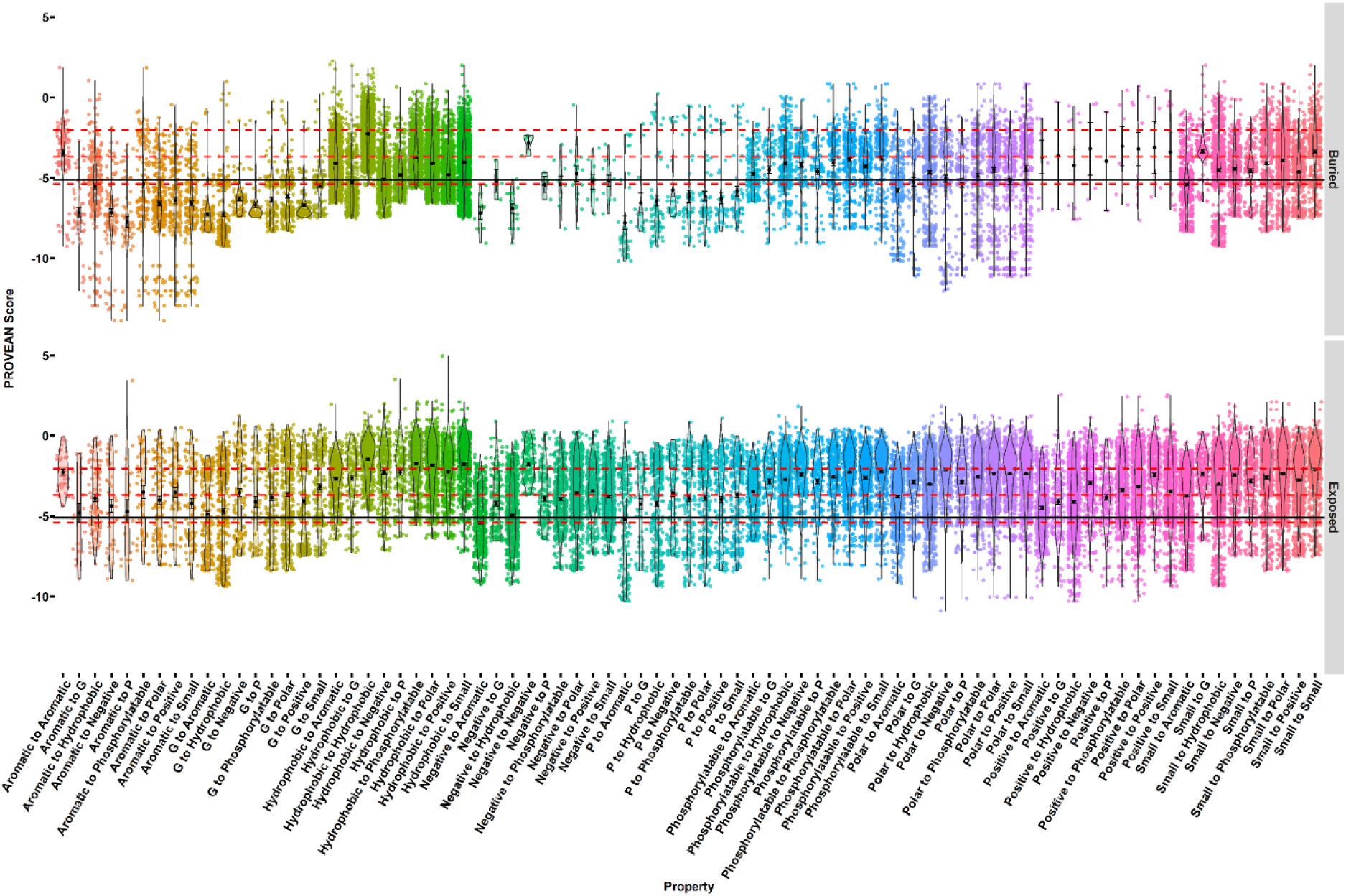
PROVEAN scores for buried and exposed residues for different mutation categories. For each category, the background dots represent the PROVEAN scores, the black lines are the distribution of the scores, black dots are the mean scores, and the bars represent the standard error. The three red horizontal lines represent the quantiles of the scores. The black horizontal line represents the mean score of the identified pathogenic mutations. G and P represent glycine and proline, respectively.

**Figure S3.**
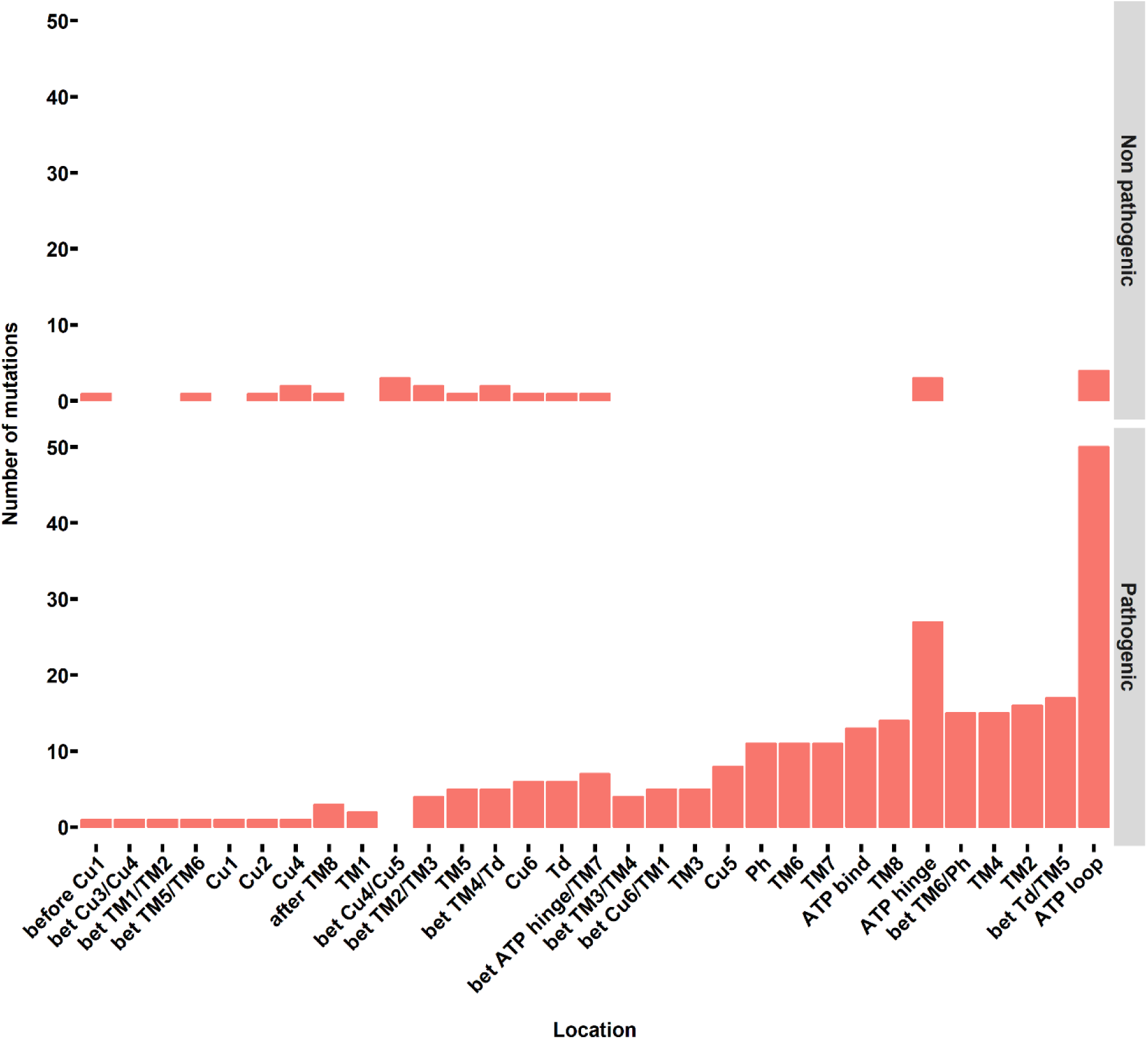
The location of the identified non-pathogenic and pathogenic mutations in the ATP7B protein.

**Figure S4.**
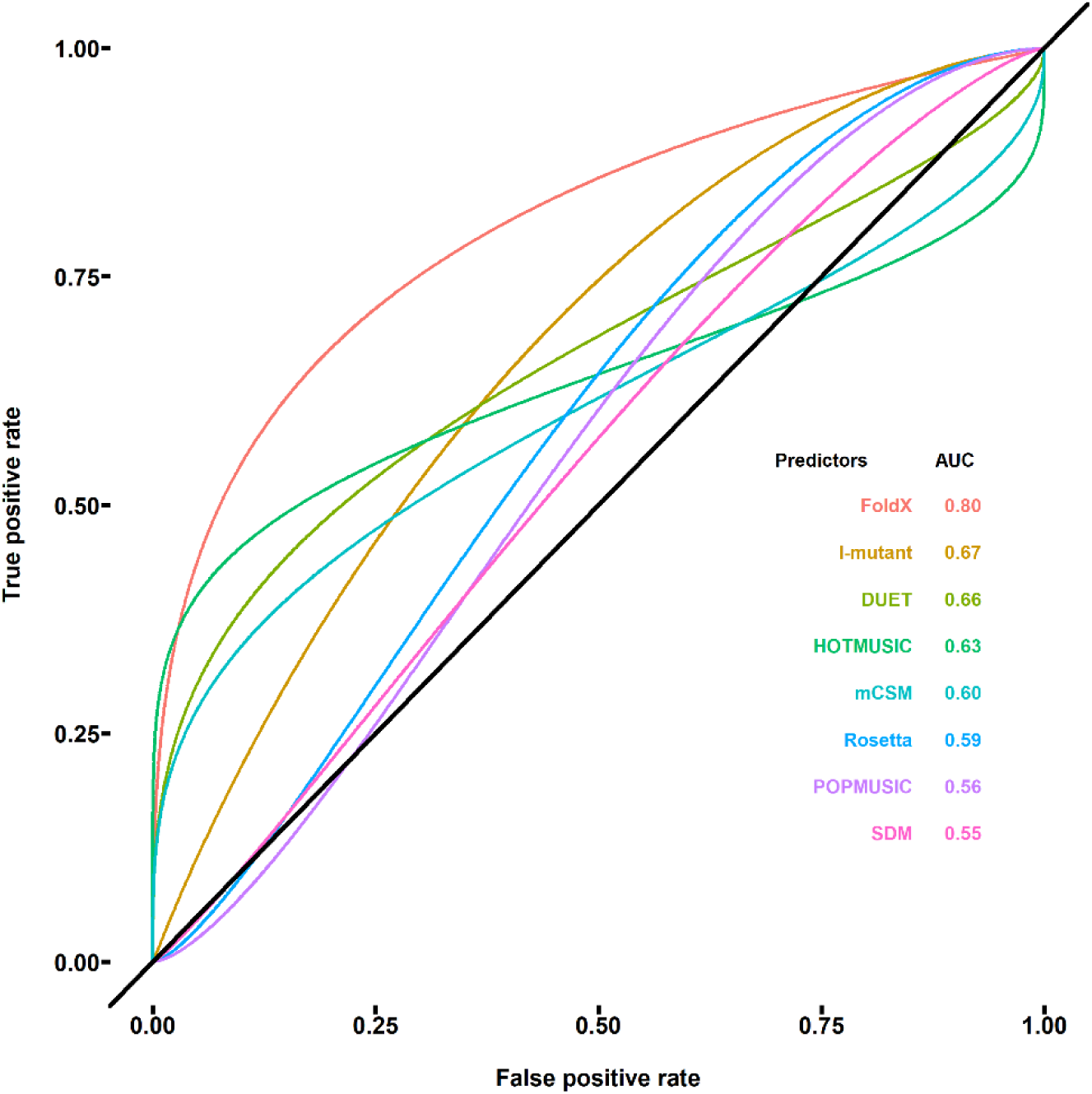
The ROC analysis for identifying the pathogenicity of mutations in the ATP binding domain using structure-based protein stability methods.

**Figure S5.**
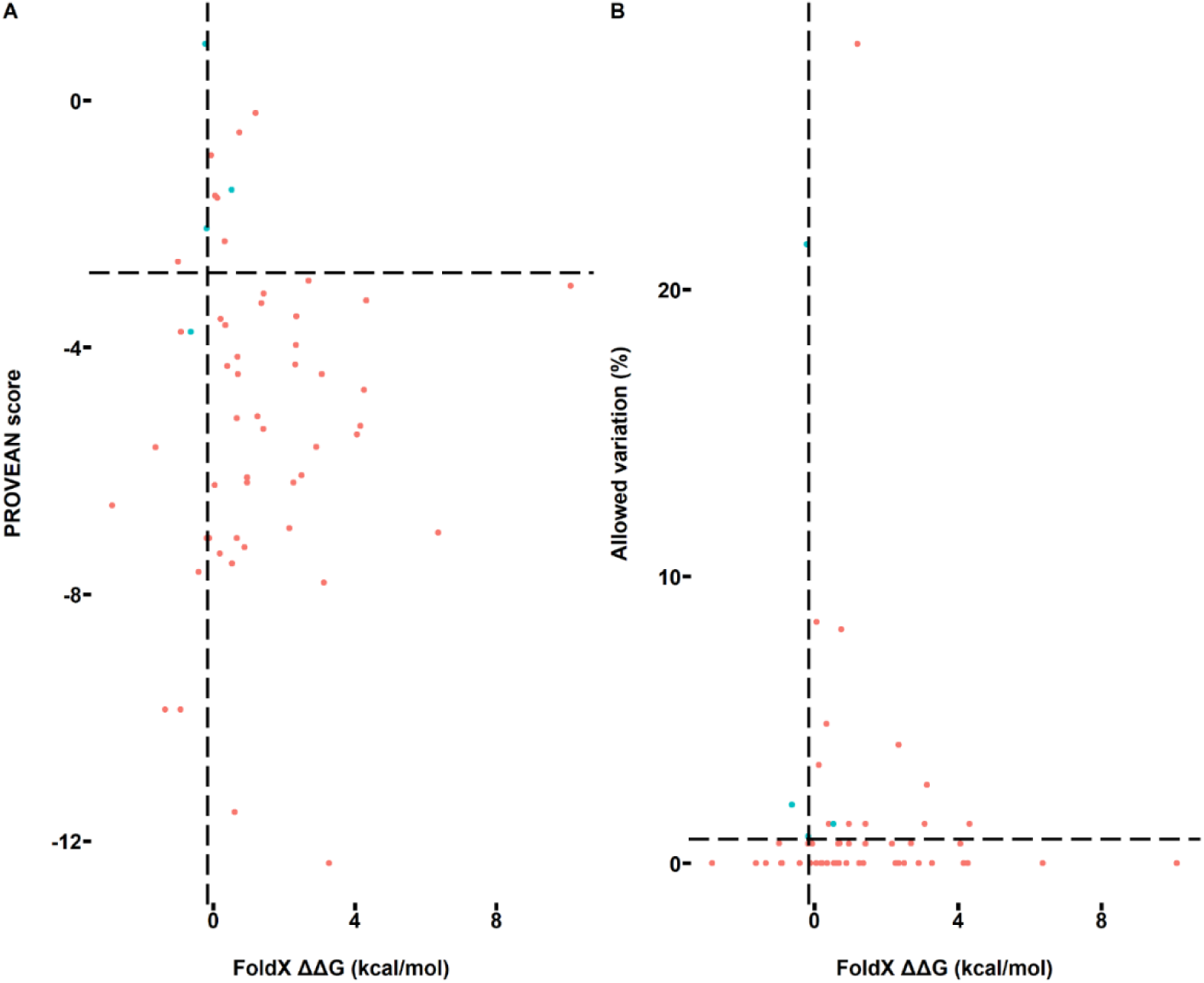
Combination of FoldX ΔΔG values and PROVEAN scores/allowed amino acid variation to create a two-dimensional representation for the identified non-pathogenic and pathogenic ATP loop binding domain (2ARF) mutations. The red and green colors represent the non-pathogenic and pathogenic mutations, respectively. The horizontal dash lines represent the PROVEAN and conservation thresholds for identifying the pathogenicity of the mutations. The vertical dash lines are the FoldX thresholds for identifying the pathogenicity of the mutations.

### Analysis using the Gao et al. 2019 data set

The following analysis were based on the recent published meta-analysis (Gao, J., Brackley, S., & Mann, J. P. (2019). The global prevalence of Wilson disease from next-generation sequencing data. Genetics in Medicine, 21(5), 1155-1163) which compiled a large amount of ATP7B variants. This data set can be downloaded from https://www.nature.com/articles/s41436-018-0309-9 (Supplementary Table 2). According to the missense mutations in Supplementary Table 2 (1315 missense mutations with clear pathogenicity: 418 pathogenic mutations and 933 non pathogenic mutations), the following analysis were performed.

**Figure S6.**
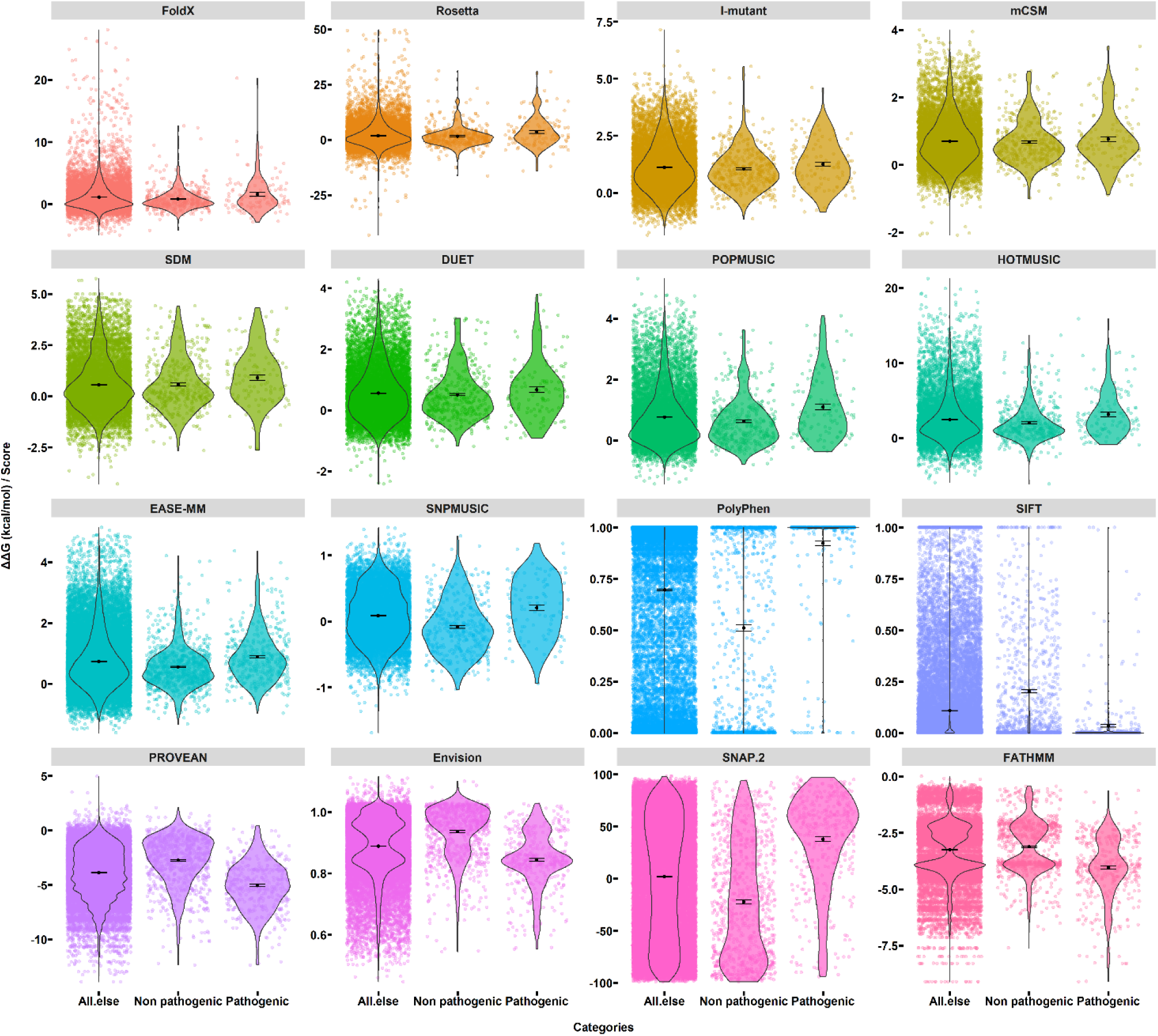
The ΔΔG values/scores for structure-based methods (FoldX, Rosetta, I-mutant, mCSM, SDM, DUET, POPMUSIC, HOTMUSIC, and SNPMUSIC) and sequence-based methods (EASE-MM, Polyphen-2, SIFT, PROVEAN, Envision, SNAP.2, and FATHMM). The background dots with jitter function represent the obtained values. The black lines represent the distribution of the obtained values. The black dots represent the mean values in each category. The error bars are the standard errors in each category.

**Table S8.**
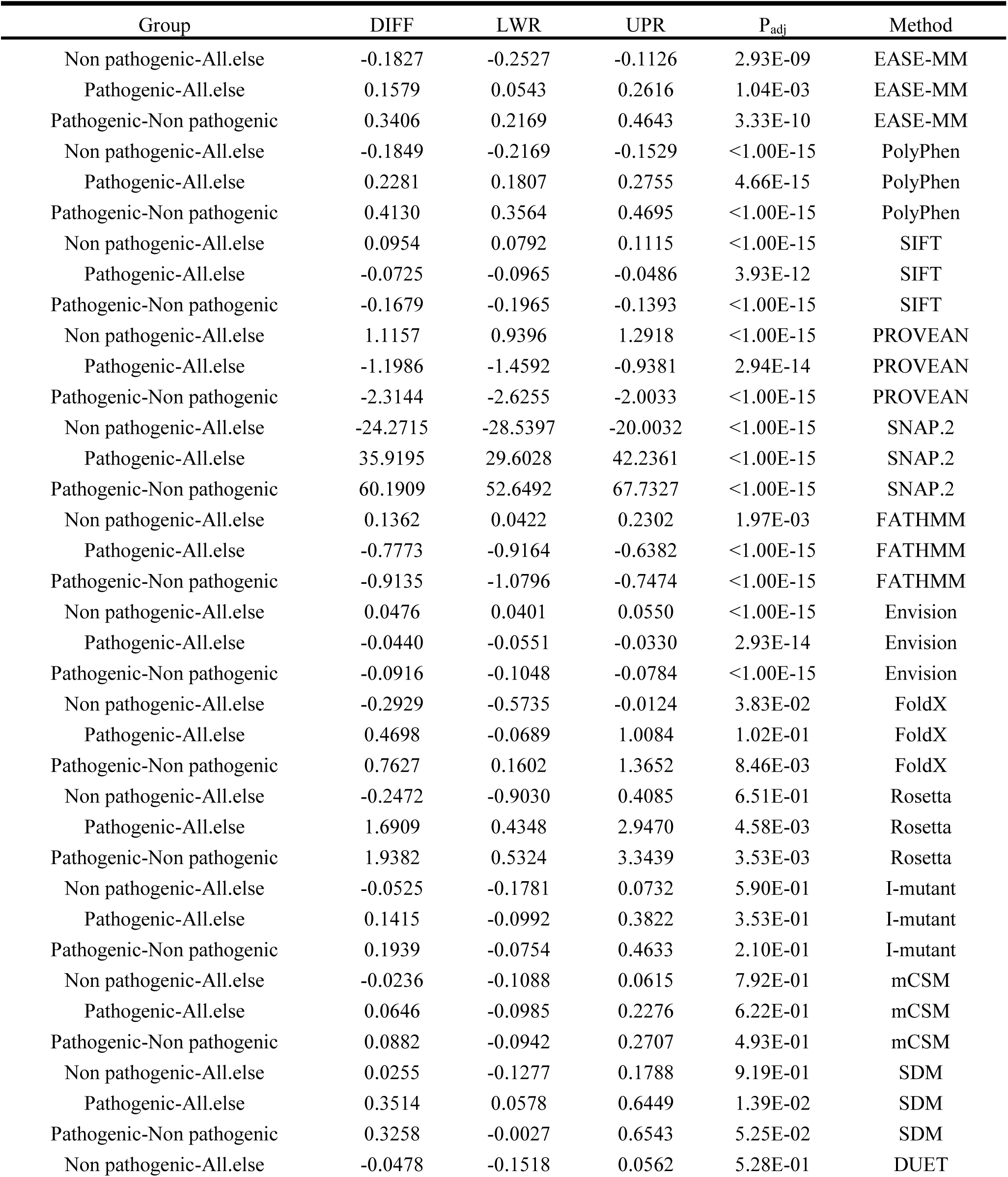
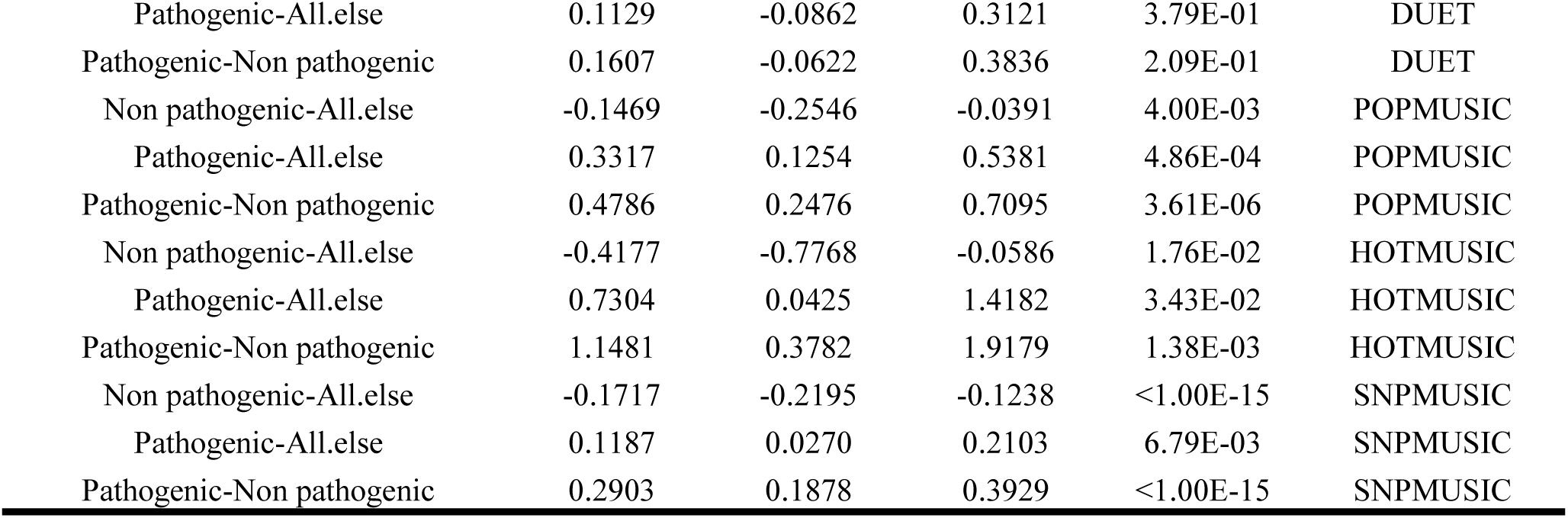
The ANOVA for all the used methods. DIFF refers to the difference between means of the two groups. LWR and UPR represent the lower and the upper end-point of the confidence interval at 95%, respectively. Padj is the adjusted p-value for multiple comparisons. Note: the signs were fixed for all the protein stability methods (ΔΔG < 0 stabilizing, ΔΔG > 0 destabilizing).

**Figure S7.**
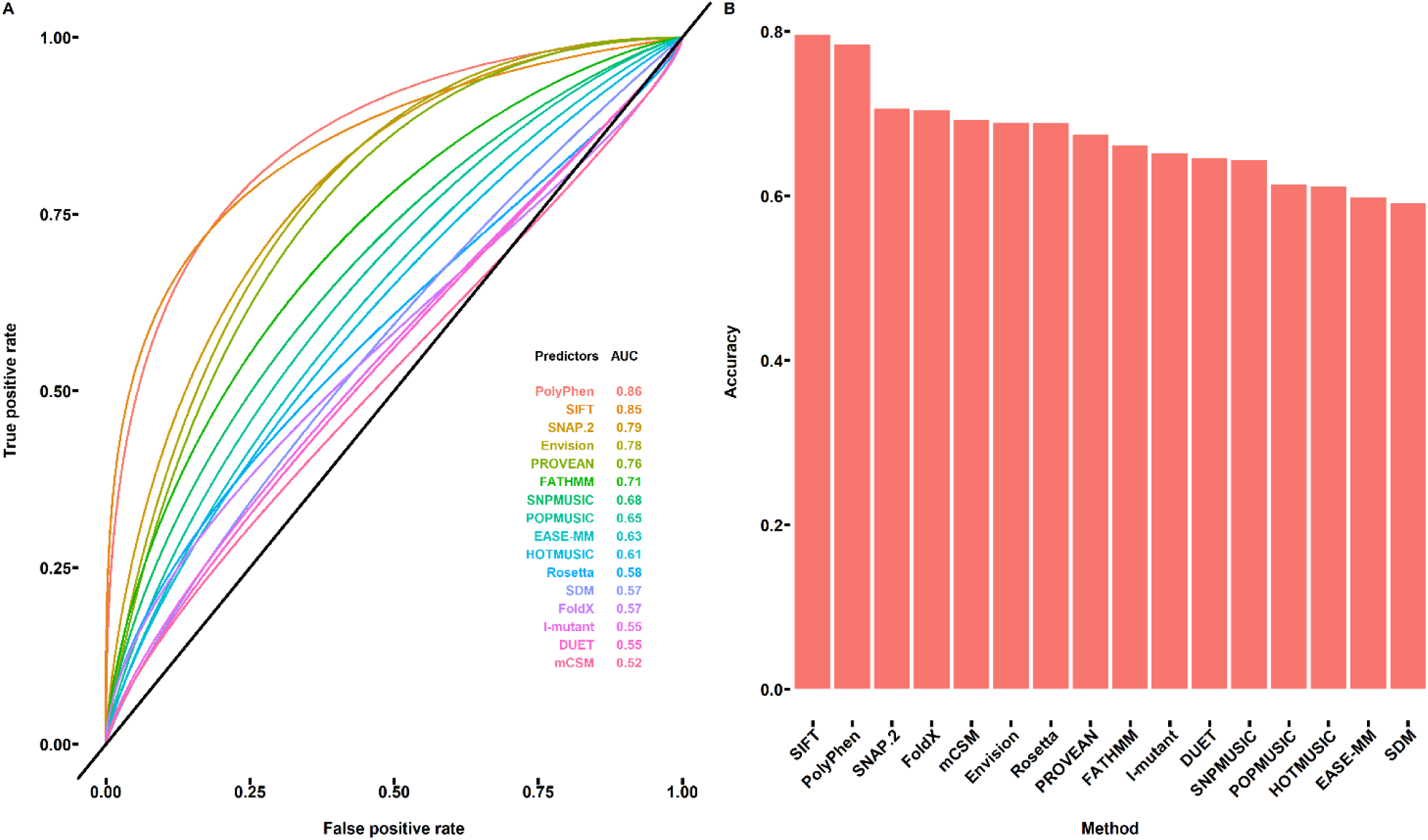
The ROC analysis. (A) The ROC plot of the benchmarked methods for identifying the pathogenicity of the ATP7B protein mutations. (B) The identification accuracy of the used methods obtained from ROC analysis.

**Figure S8.**
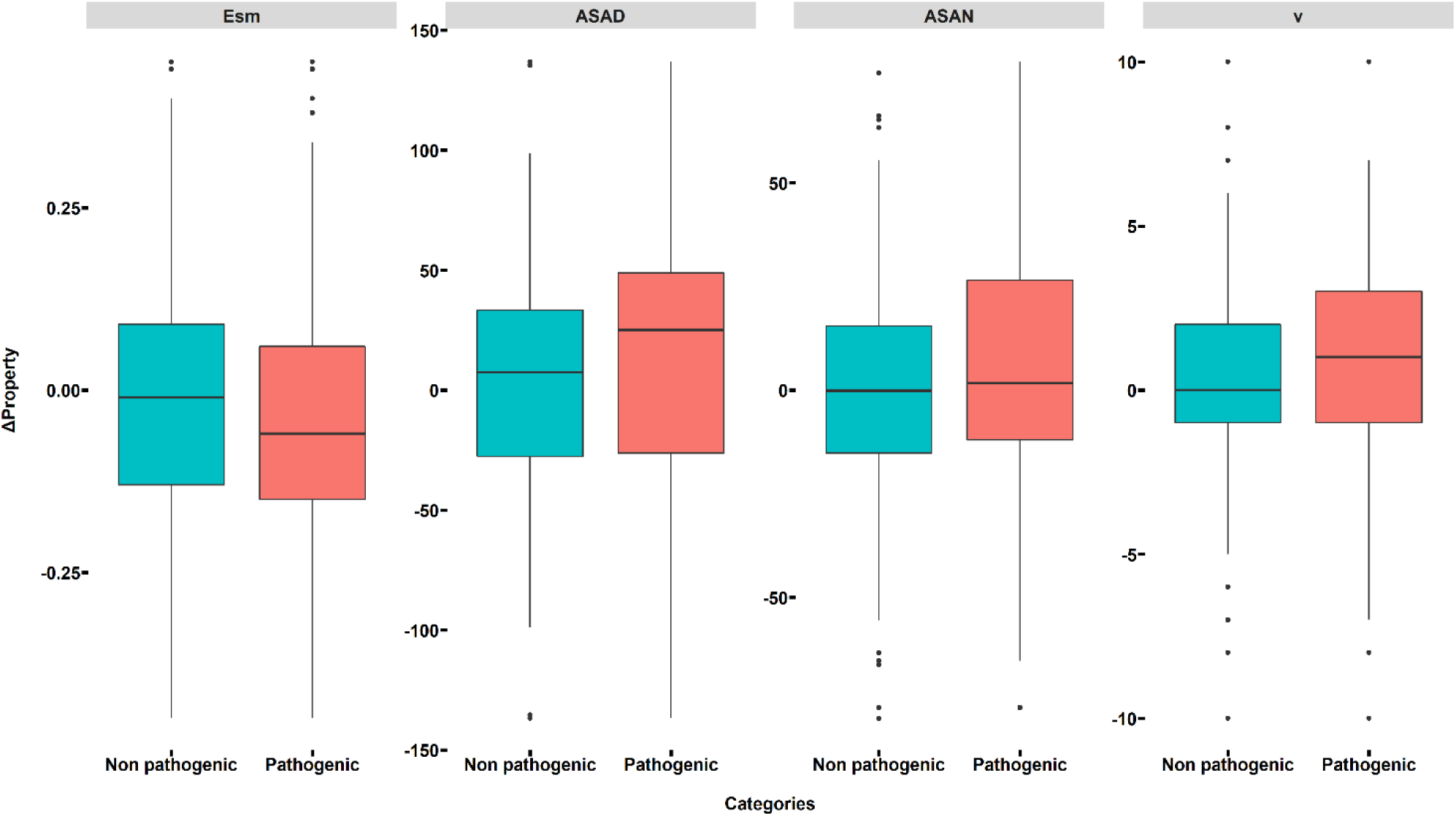
Distribution of amino acid property changes (p < 0.001) for non-pathogenic and pathogenic ATP7B mutations. Thick bars indicate the median in each category; the edges of the color-filled rectangles represent the 25th and 75th percentiles. The black dots represent the outliers of the range covered by the black bars. Esm: short- and medium-range non-bonded energy; ASAD and ASAN are solvent-accessible surface area for denatured and native, respectively. v: volume (number of non hydrogen sidechain atoms).

**Table S9.** The ANOVA for amino acid property changes. DIFF refers to the difference between means of the two groups. LWR and UPR represent the lower and the upper end-point of the confidence interval at 95%, respectively. P_adj_ is the adjusted p-value for multiple comparisons.

| Group | DIFF | LWR | UPR | $P_{adj}$ | Method |
| --- | --- | --- | --- | --- | --- |
| Pathogenic-Non pathogenic | 0.0971 | -0.4146 | 0.6088 | 0.709707 | K |
| Pathogenic-Non pathogenic | 0.2074 | 0.0435 | 0.3713 | 0.013178 | Ht |
| Pathogenic-Non pathogenic | -0.1066 | -0.2885 | 0.0753 | 0.250527 | Hs |
| Pathogenic-Non pathogenic | 2.4720 | -0.6673 | 5.6114 | 0.122641 | P |
| Pathogenic-Non pathogenic | 0.3385 | 0.0726 | 0.6044 | 0.012631 | pHi |
| Pathogenic-Non pathogenic | -0.0590 | -0.1048 | -0.0131 | 0.011741 | pK |
| Pathogenic-Non pathogenic | 7.3763 | 2.9854 | 11.7671 | 0.001008 | Mw |
| Pathogenic-Non pathogenic | 0.9627 | 0.2314 | 1.6939 | 0.009914 | B1 |
| Pathogenic-Non pathogenic | 0.3326 | -0.3356 | 1.0008 | 0.328960 | Rf |
| Pathogenic-Non pathogenic | 1.5566 | 0.2371 | 2.8760 | 0.020801 | u |
| Pathogenic-Non pathogenic | -0.2122 | -0.3603 | -0.0641 | 0.005021 | Hnc |
| Pathogenic-Non pathogenic | -0.0346 | -0.0549 | -0.0143 | 0.000834 | Esm |
| Pathogenic-Non pathogenic | -0.0051 | -0.0226 | 0.0124 | 0.566588 | El |
| Pathogenic-Non pathogenic | -0.0407 | -0.0665 | -0.0150 | 0.001940 | Et |
| Pathogenic-Non pathogenic | -0.0078 | -0.0543 | 0.0386 | 0.740869 | Pa |
| Pathogenic-Non pathogenic | -0.0061 | -0.0601 | 0.0478 | 0.823896 | Pb |
| Pathogenic-Non pathogenic | 0.0010 | -0.0542 | 0.0562 | 0.970642 | Pt |
| Pathogenic-Non pathogenic | 0.0096 | -0.0390 | 0.0582 | 0.698324 | Pc |
| Pathogenic-Non pathogenic | 2.5851 | 0.7662 | 4.4040 | 0.005377 | Ca |
| Pathogenic-Non pathogenic | 0.0012 | -0.0197 | 0.0222 | 0.907572 | F |
| Pathogenic-Non pathogenic | -0.0306 | -0.0535 | -0.0078 | 0.008706 | Br |
| Pathogenic-Non pathogenic | -0.0582 | -0.3093 | 0.1929 | 0.649236 | Ra |
| Pathogenic-Non pathogenic | -0.1009 | -0.2239 | 0.0221 | 0.107798 | Ns |
| Pathogenic-Non pathogenic | -0.0777 | -0.1578 | 0.0024 | 0.057148 | an |
| Pathogenic-Non pathogenic | 0.0257 | -0.0662 | 0.1175 | 0.583769 | ac |
| Pathogenic-Non pathogenic | 0.0784 | -0.0045 | 0.1613 | 0.063868 | am |
| Pathogenic-Non pathogenic | 5.8075 | 2.0401 | 9.5750 | 0.002542 | V0 |
| Pathogenic-Non pathogenic | -0.0252 | -0.0726 | 0.0221 | 0.296381 | Nm |
| Pathogenic-Non pathogenic | -0.0660 | -0.1798 | 0.0479 | 0.256058 | NI |
| Pathogenic-Non pathogenic | -0.0416 | -0.1917 | 0.1085 | 0.586797 | Hgm |
| Pathogenic-Non pathogenic | 10.2761 | 4.5671 | 15.9851 | 0.000428 | ASAD |
| Pathogenic-Non pathogenic | 5.7856 | 2.5516 | 9.0196 | 0.000464 | ASAN |
| Pathogenic-Non pathogenic | 4.4755 | -0.1429 | 9.0939 | 0.057512 | ΔASA |
| Pathogenic-Non pathogenic | -0.2998 | -0.5907 | -0.0089 | 0.043423 | ΔGh |
| Pathogenic-Non pathogenic | -0.4977 | -1.1818 | 0.1863 | 0.153700 | GhD |
| Pathogenic-Non pathogenic | -0.2421 | -0.5615 | 0.0772 | 0.137153 | GhN |
| Pathogenic-Non pathogenic | -0.5372 | -0.9282 | -0.1463 | 0.007114 | ΔHh |
| Pathogenic-Non pathogenic | 0.2374 | 0.0660 | 0.4088 | 0.006662 | TΔSh |
| Pathogenic-Non pathogenic | 1.1716 | -0.5339 | 2.8771 | 0.178014 | ΔCh |
| Pathogenic-Non pathogenic | 0.3087 | 0.0358 | 0.5816 | 0.026628 | $\Delta G_c$ |
| Pathogenic-Non pathogenic | 0.4681 | 0.0634 | 0.8729 | 0.023422 | $\Delta H_c$ |
| Pathogenic-Non pathogenic | -0.1603 | -0.4437 | 0.1231 | 0.267260 | $T\Delta S_c$ |
| Pathogenic-Non pathogenic | 0.0092 | -0.0806 | 0.0990 | 0.840148 | $\Delta G$ |
| Pathogenic-Non pathogenic | -0.0702 | -0.4126 | 0.2723 | 0.687824 | $\Delta H$ |
| Pathogenic-Non pathogenic | 0.0800 | -0.1830 | 0.3431 | 0.550802 | $T\Delta S$ |
| Pathogenic-Non pathogenic | 0.5588 | 0.2330 | 0.8846 | 0.000787 | v |
| Pathogenic-Non pathogenic | 0.0860 | -0.1653 | 0.3372 | 0.502183 | s |
| Pathogenic-Non pathogenic | 0.2316 | 0.0123 | 0.4510 | 0.038479 | f |

**Figure S9.**
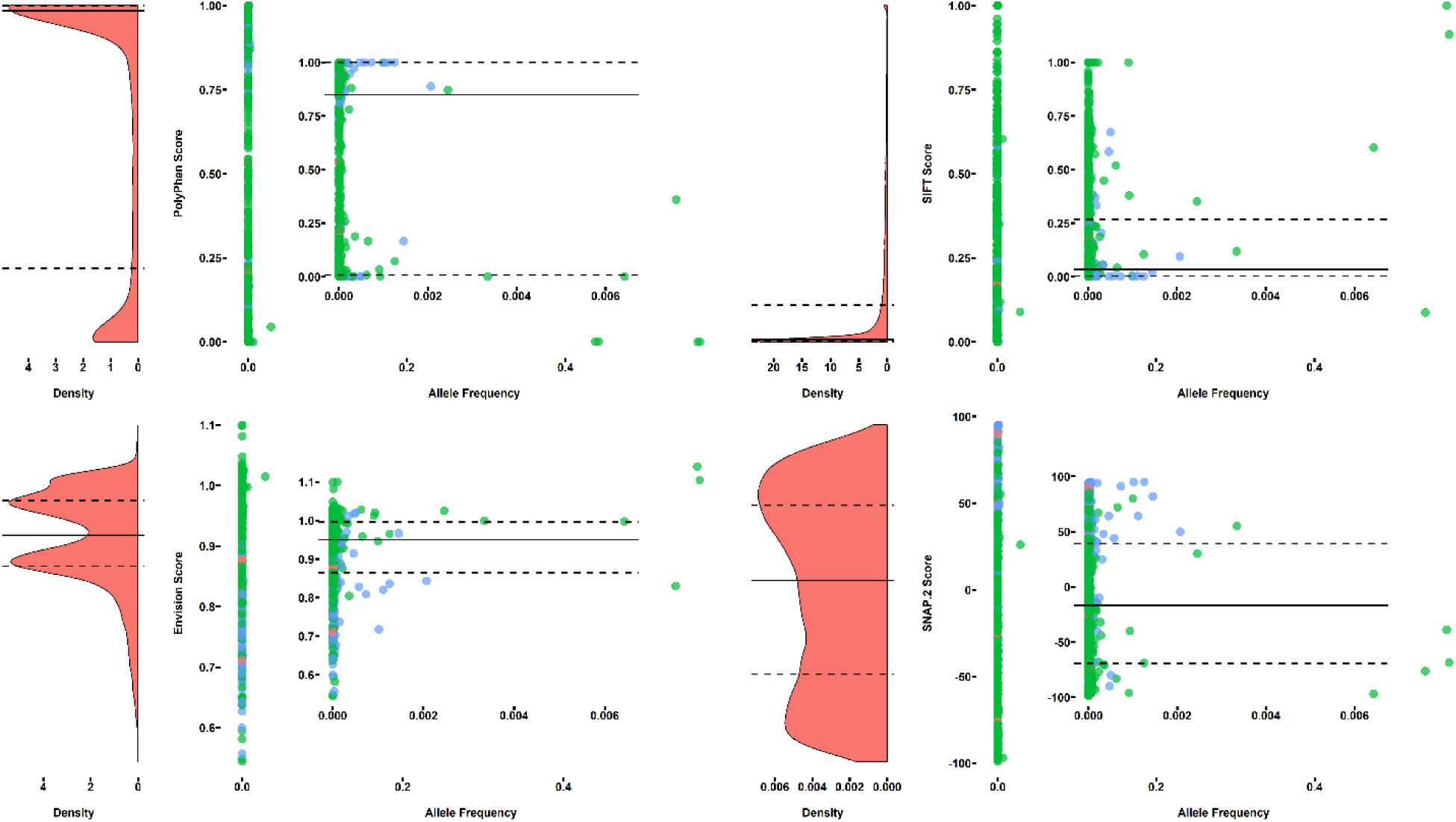
The correlation between PolyPhen, SIFT, Envision, and SNAP.2 scores and allele frequency. The plots on the left side in each category are the density plots of the scores. The inset plots on the right side are the zoom-in correlation plots with the allele frequency lower than 0.009. The red, green and blue dots represent the ATP7B protein mutations with unknown pathogenicity, non-pathogenic mutations, and pathogenic mutations, respectively.

**Figure S10.**
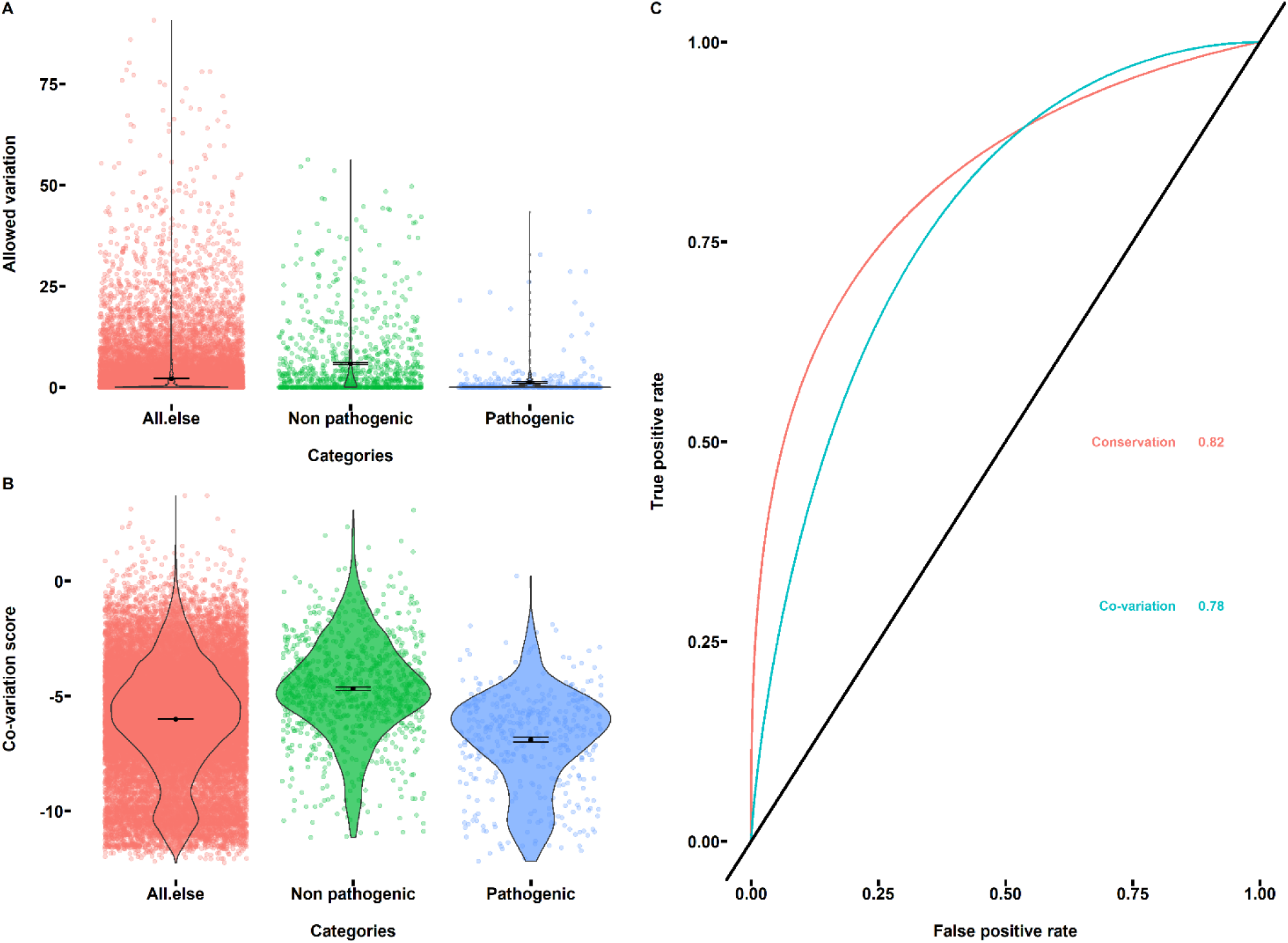
Conservation analysis and co-variation analysis used to identify pathogenic ATP7B mutations. (A) The allowed amino acid variation at each position and variation distributions for different categories. The dots in the background represent the obtained allowed variation; the black lines represent the variation distribution. The black dots represent the mean values in each category, and the error bars represent the standard errors in each category. (B) The co-variation scores obtained from GREMLIN and score distributions for different categories. (C) The ROC plot for conservation analysis and co-variation analysis with the final AUC value labeled.

**Figure S11.**
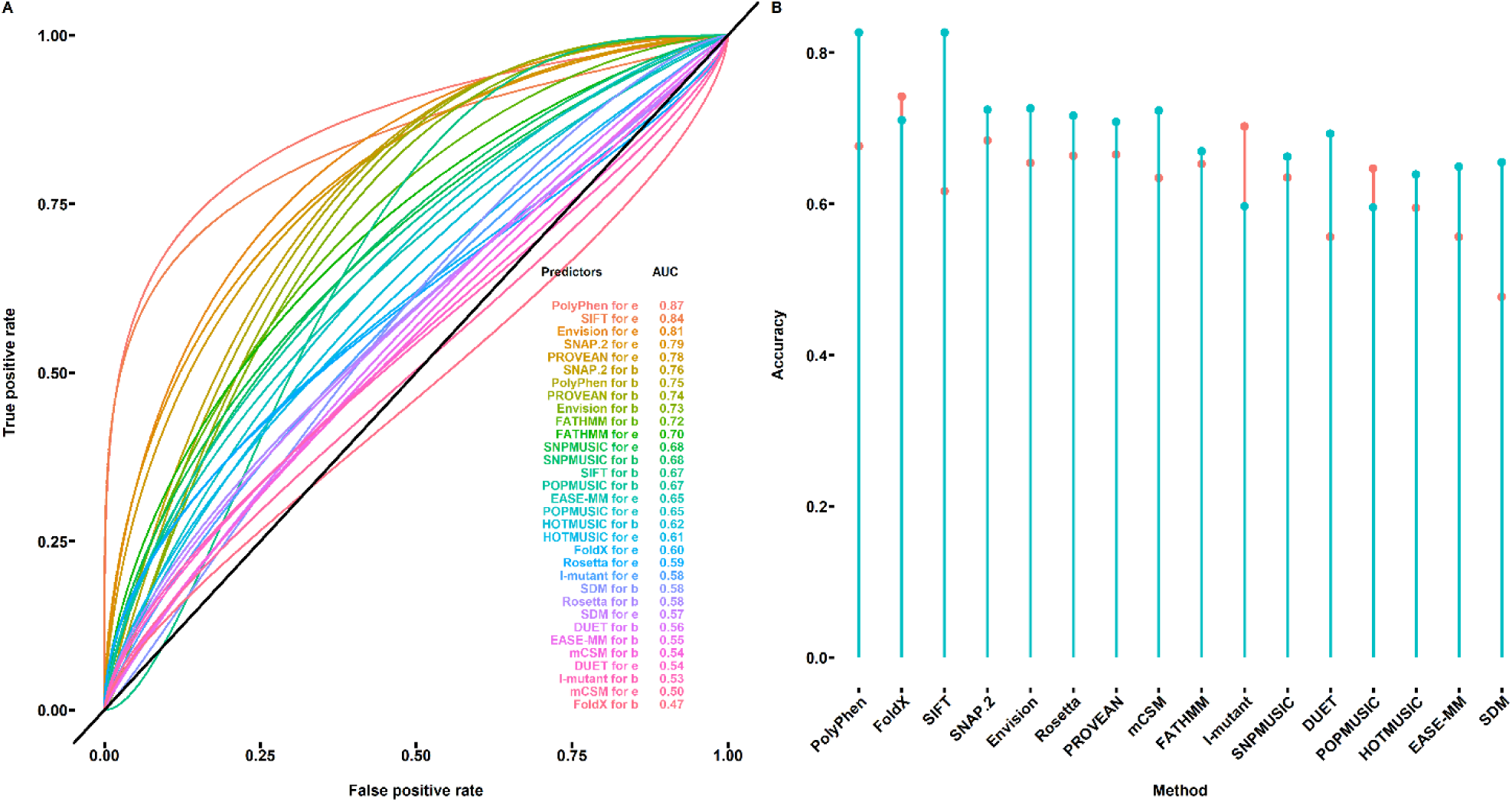
ROC analysis for buried and exposed residues. (A) ROC plot of the used structure-based and sequence-based methods applied to predict the pathogenicity of mutations in buried (b) and exposed (e) sites. (B) Identification accuracy of the used methods obtained from ROC analysis. The red and green colors represent the accuracy for buried and exposed residues, respectively.

**Figure S12.**
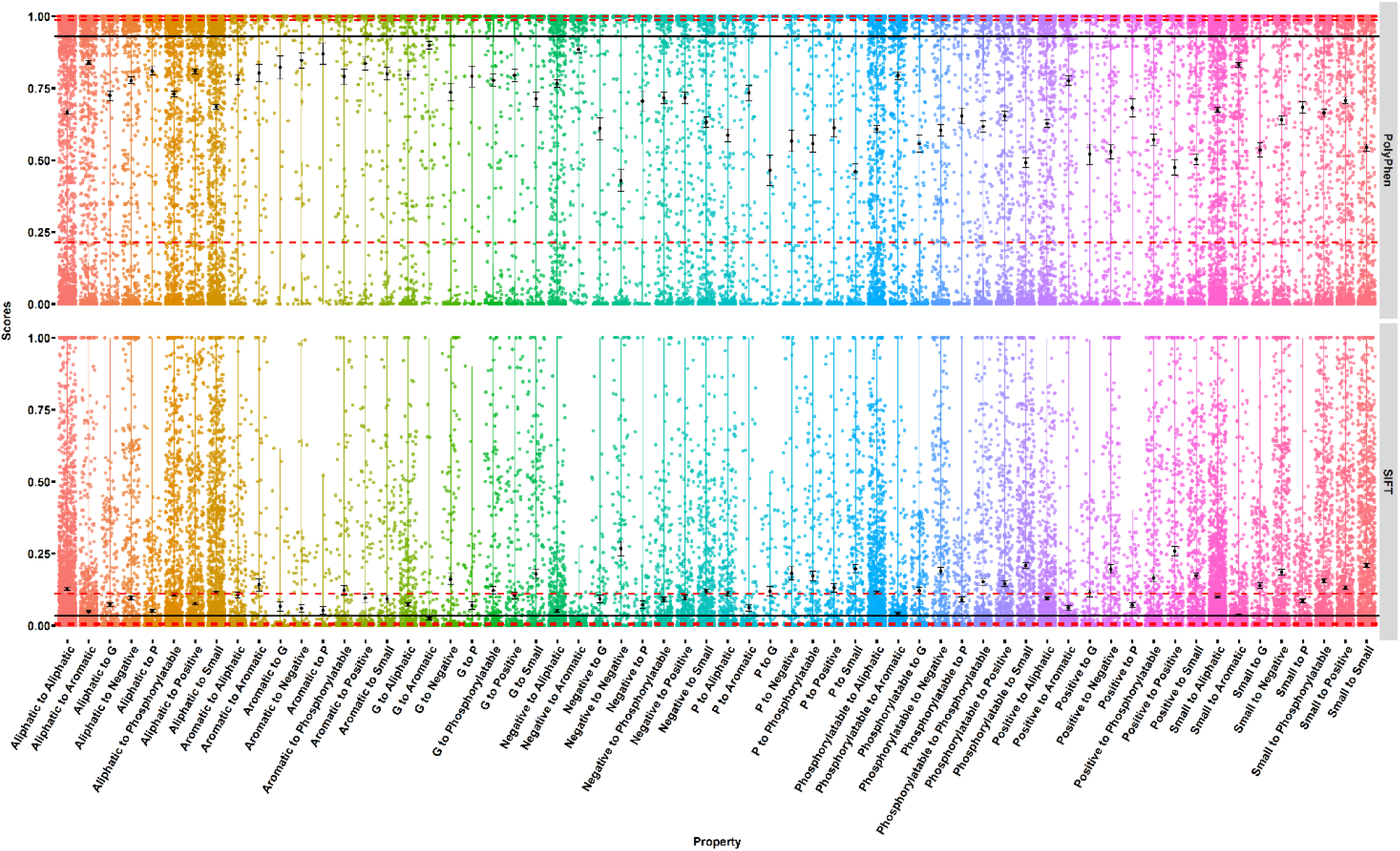
PolyPhen and SIFT scores for different mutation categories. For each category, the background dots represent the PolyPhen and SIFT scores, the black lines are the distribution of the scores, black dots are the mean scores, and the bars represent the standard error. The three red horizontal lines represent the quantiles of the scores. The black horizontal line represents the mean score of the identified pathogenic mutations. G and P represent glycine and proline, respectively.

